# *sisterless A* is required for activation of *Sex lethal* in the *Drosophila* germline

**DOI:** 10.1101/2019.12.17.880070

**Authors:** Raghav Goyal, Ellen Baxter, Mark Van Doren

## Abstract

Both somatic cells and germ cells must establish their correct sexual identity for proper gametogenesis. In *Drosophila,* sex determination in somatic cells is controlled by the switch gene *Sex lethal* (*Sxl*), which is activated in females by the presence of two X chromosomes. Though germline sex determination is much less well understood, *Sxl* is also essential for the female identity in germ cells. Loss of *Sxl* function in the germline results in ovarian germline tumors, a characteristic of male germ cells developing in a female soma. Further, *Sxl* expression is sufficient for XY (male) germ cells to produce eggs when transplanted into XX (female) somatic gonads. As in the soma, the presence of two X chromosomes activates *Sxl* in the germline, but the mechanism for “counting” X chromosomes in the germline is thought to be different from the soma. Here we have explored this mechanism at both *cis*- and *trans-* levels. Our data support the model that the *Sxl* “establishment” promoter (*SxlPE*) is activated in a female-specific manner in the germline, as in the soma, but that the timing of *SxlPE* activation, and the DNA elements that regulate *SxlPE,* are different in the germline. Nevertheless, we find that the X chromosome gene *sisterless A (sisA),* which helps activate *Sxl* in the soma, is also essential for *Sxl* activation in the germline. Loss of *sisA* leads causes of Sxl expression in the germline, and to ovarian tumors and germline loss. These defects can be rescued by Sxl expression, demonstrating that *sisA* lies upstream of *Sxl* in germline sex determination. We conclude that *sisA* acts as an X chromosome counting element in both the soma and the germline, but that additional factors regulating female-specific expression of *Sxl* in the germline remain to be discovered.

**AUTHOR SUMMARY:** The production of sperm and eggs requires proper sexual identity to be established in both somatic cells and the germ cells, which ultimately produce the gametes. While somatic sex determination has been well studied in a number of organisms, how germ cells establish their sexual identity is much less well understood. In Drosophila, the RNA binding protein Sex lethal (Sxl) is essential for female sexual identity in both the soma and the germline, but its regulation in the germline is thought to be different than in the soma. Here we explore how *Sxl* is activated in the germline. We find that the germline uses a different set of DNA elements to control activation of the key sex-specific *Sxl* promoter. Nonetheless, one of the activators of *Sxl* in the soma, the transcription factor Sisterless A (SisA), also acts to activate *Sxl* in the germline. Our data indicate that, while SisA acts as a common activator in both the soma and germline, additional, germline-specific *Sxl* activators remain to be discovered.

## INTRODUCTION

Sex determination influences the development of many different tissues, but is particularly important in the germline to control the production of either sperm or eggs. While in some species somatic sex is sufficient to determine germline sex via inductive signaling, in flies and humans, the sex chromosome karyotype also plays a role intrinsically in the germline. Since a failure to match germline and somatic sex leads to defects in gametogenesis, understanding how intrinsic sex determination is regulated in the germline is important for our understanding of gonad development, reproductive biology, and human health.

Sex determination in *Drosophila* is under the control of the switch gene *Sex lethal* (*Sxl*) which is activated in females by the presence of two X chromosomes [1–11]. *Sxl* is both necessary and sufficient for the female sexual identity of somatic cells [3,6,12–19]. Expression of *Sxl* in the soma is regulated by two different promoters [20,21]. The *Sxl* “establishment promoter” (*SxlPE*) is sex-specific and is activated by the presence of two X chromosomes [2]. Default splicing of the *SxlPE* transcript produces a pulse of ‘Early’ Sxl protein starting at embryonic nuclear cycle 12. Later, at nuclear cycle 14, dosage compensation equalizes X chromosome expression in males and females and so X chromosomes can no longer be counted. Thus, there is a transition to the “maintenance promoter,” *SxlPM*, which is activated in both sexes [20,21]. The transcript from *SxlPM* requires alternative splicing regulated by the early Sxl protein (only made in females) in order to produce more ‘Late’ Sxl protein, creating an auto-regulatory loop that maintains Sxl expression in females [3]. The initial, female-specific activation of *SxlPE* in the soma is in direct response to a diploid dose of X-linked Signaling Elements (XSEs) – genes on the X chromosome that activate *SxlPE* when present in two copies, but not one. The somatic XSEs include the transcription factors Sisterless A (SisA), Sisterless B (SisB) and Runt, along with the JAK/STAT ligand Unpaired (Upd) [9,22–30].

Unlike primary sex determination in the soma, which is autonomously dependent on a cell’s sex chromosome genotype, sex determination in the *Drosophila* germline is regulated by non-autonomous signals from the soma in addition to the sex chromosome make-up of the germline (reviewed in [31]). In order for proper gametogenesis to occur, the “sex” of the germ cells has to match the sex of the soma. For example, XX germ cells developing in a male soma result in a testis with a severely atrophic germline [32–45], while XY germ cells developing in a female soma produce an ovary with a tumorous germline (“ovarian tumor”) and a complete failure to produce eggs [32,46–48]. Similar to the soma, *Sxl* is both necessary and sufficient for determining the autonomous component of germline sexual identity. *Sxl* loss-of-function in the germline results in ovarian germline tumors, similar to male germ cells developing in a female soma [32,33,37,39,42,46,48,49]. Further, XY (male) germ cells expressing *Sxl* are able to produce eggs when transplanted into XX (female) somatic gonads, demonstrating that *Sxl* is sufficient for female sexual identity in the germline as long as the surrounding soma is female [50]. Female-specific Sxl expression in the germline is dependent on the X chromosome dose: XX germ cells express Sxl while XY germ cells do not [32,39]. Evidence also indicates that *SxlPE* is important for female-specific expression of Sxl in the germline. Transcript data from early germ cells show a *Sxl* RNA that matches the *SxlPE* transcript in the soma [50], and data indicate that the positive auto-regulation of *Sxl* expression that occurs in the soma also acts in the germline [49]. However, the mechanism for activating *Sxl* expression in the germline appears to be different from the soma [33,46,50–54]. When germ cells simultaneously heterozygous for *Sxl* and the somatic XSEs *sisA*, *sisB*, and *runt*, were transplanted into wildtype female embryos, they were still able to undergo oogenesis indicating that the germ cells were not masculinized, even though this same genotype is sufficient to masculinize somatic cells [52]. Further, germ cells homozygous for mutations in *sisB*, or maternally deficient for the *sisB* co-factor *daughterless*, can also make normal eggs [33,46]. Thus, different XSEs, or at least a different combination of XSEs, is required to activate Sxl expression in the germline.

Here we utilize a combination of approaches to investigate the female-specific activation of *Sxl* in the germline. Using RNA FISH and CRISPR-tagging of specific Sxl isoforms, we show that the timing of *SxlPE* activation relative to *SxlPM* is different in the germline than in the soma. Using promoter/enhancer reporter constructs, we demonstrate that the regulatory sequences required for female-specific activation of *SxlPE* in the germline are different from those required in the soma. Lastly, we show that the somatic XSE, *sisA,* is also required for female-specific expression of Sxl in the germline. Together our data support a model in which *sisA* acts as a germline XSE, but that additional factors, different from the somatic XSEs, are also important for *Sxl* activation in the female germline.

## RESULTS

### Analysis of *Sex lethal* transcription in PGCs

Since the activation of *SxlPE* appears to be the key sex-specific decision in germline sex determination, similar to what is observed in the soma, we first decided to examine when and how *SxlPE* is activated in the germline. We hypothesized that, as in the soma, *SxlPE* would be activated prior to *SxlPM*. To study this, we performed RNA fluorescent *in situ* hybridization (FISH) using oligopaints to examine nascent RNA (nRNA) being transcribed from *SxlPE* and *SxlPM* at the *Sxl* locus. As *SxlPM* is upstream of *SxlPE*, we generated one set of probes that exclusively targets *SxlPM*-derived transcripts (*PM*-probe, Cy5, Green), and another that targets the common transcript from both *SxlPE* and *SxlPM* (*Com*-probe, Cy3, Red) (Fig. S1A). Thus, if *SxlPE* alone is active, we should observe signal from the *Com*-probe (Red) but not the *PM*-probe (Green), while if *SxlPM* is active we should observe signal from both probes (and cannot make a conclusion about *SxlPE*).

In the soma, female-specific activation of *SxlPE* occurs prior to expression of *SxlPM*. Consistent with this, in the female soma we initially observed RNA FISH signals as fluorescent nuclear foci with the *Com*-probe but not the *PM*-probe, indicating that only *SxlPE* was activated (Fig. S1C-S1C’’). Subsequently, we observed signals from both probes, indicating that *SxlPM* had been activated (Data not shown). Conversely, in the male soma, we always observed a signal from both the *PM-* and *Com*-probes simultaneously (Fig. S1B-S1B’’), indicating that that *SxlPE* is not expressed (or does not precede *SxlPM* activation). We observed two *Sxl* foci per nucleus in females and only one focus in males, which was expected as *Sxl* is an X chromosome locus. These data are consistent with the previous understanding of *Sxl* activation in the soma - that *SxlPE* is on initially only in females followed by *SxlPM* activation in both sexes.

In contrast, in both female and male germ cells, we only observed RNA FISH signals from both the *PM*-probes and the *Com-*probes simultaneously, and never from the *Com*-probe alone (Fig. 1A-1B’’). This indicates that *SxlPE* is not activated before *SxlPM* in the germline. (Note: we only observed a single focus in both XX and XY germ cells suggesting that the X chromosomes may be paired in the germline.) The simultaneous expression of *PM*-probes and *Com*-probes occurred in a subset of female germ cells at stage 5 (11.1%), and increased to 100 % of germ cells by stage 11 (Fig. 1C). Similarly, both probe sets were expressed simultaneously in a subset of male germ cells, starting at stage 5 (7%) and increased to 100% of germ cells by stage 11 (Fig. 1D). The gradual increase in the number of cells expressing *SxlPM* transcripts is similar to what is observed in the soma [21]. We conclude that, unlike in female somatic cells, *SxlPE* does not precede *SxlPM* activity in the germline, but instead is likely activated either simultaneously with, or later than, *SxlPM*. This is still consistent with a model where ‘Early Sxl’ protein produced from default splicing of the *SxlPE*-derived transcript is required for productive RNA splicing of the *SxlPM*-derived transcript, and thus for the subsequent production of ‘Late Sxl’ protein; even if *SxlPM* is active prior to *SxlPE* activation, these transcripts would not be able to produce Sxl protein until *SxlPE* is activated.

**Figure 1:**
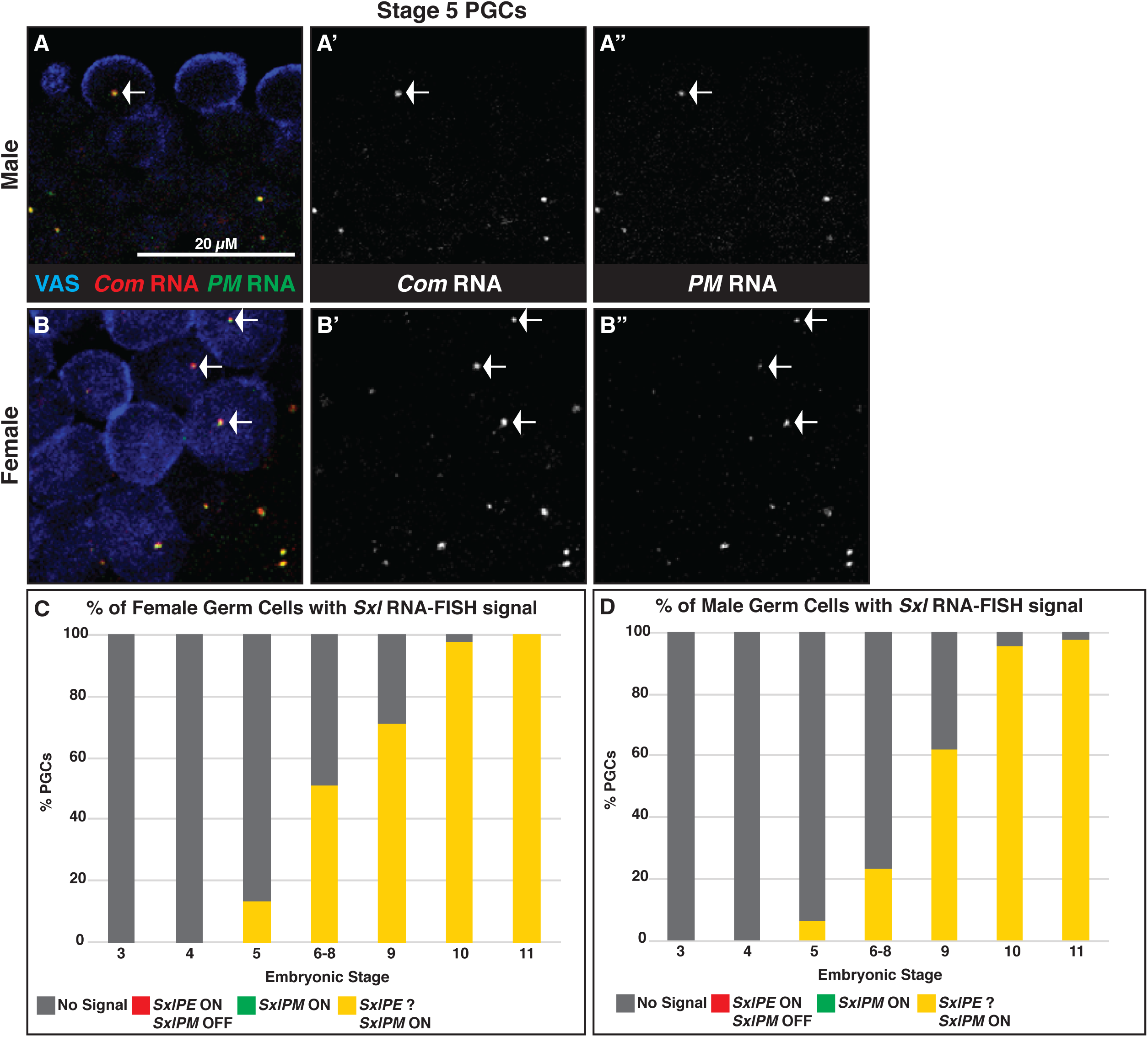
*SxlPE* activity does not precede *SxlPM* in the germline. A-B) RNA-FISH (+ Immunofluorescence) against the transcript from *SxlPM* only (*PM* RNA) versus the common transcript from *SxlPE* and *SxlPM* (*PE+PM* RNA) in embryonic stage 5 primordial germ cells (PGCs). A) Male PGCs with both *PE+PM* and *PM*-probe signals. B) Female PGCs with both *PE+PM* and *PM*-probe signals. Arrows mark fluorescent nuclear foci of RNA-FISH signals. Note that a single focus is observed in female PGCs suggesting that X chromosomes may be paired in the germ cells. VAS stains germ cells. C-D) Graphs showing percentage of PGCs with observable RNA-FISH probe signals between stages 3 and 11 of embryogenesis.

### *SxlPE* activity in the germline requires different *cis*-regulatory elements than in the soma

We next wanted to identify if and when *SxlPE* is activated in the germline, and also to compare the *cis-*regulatory logic of *SxlPE* activation in the germline to that in the soma. To do this we used transcriptional reporter constructs specific for *SxlPE.* We first tested a *SxlPE*-EGFP transcriptional reporter that contains the 1.5kb somatic enhancer immediately upstream of *SxlPE* (*SxlPE*-1.5kb), which had previously been reported to recapitulate both somatic and germline *SxlPE* activity [50,74]. Contrary to previous reports, we did not observe EGFP expression from *SxlPE*-1.5kb in developing PGCs, even though *SxlPE-*1.5kb showed sex-specific EGFP expression in somatic cells (Data not shown). Thus, we conclude that the *cis-*regulatory region sufficient for sex-specific expression of *SxlPE* in the soma is insufficient for *SxlPE* expression in the germline. We next extended the *SxlPE* transcriptional reporter to contain the entire 5.2kb genomic sequence upstream of *SxlPE* (but excluding *SxlPM*) (*SxlPE-*5.2kb, Fig. S2A). Once again, even though we observed female-specific nuclear EGFP expression from the *SxlPE-*5.2kb reporter in somatic cells, we found no evidence of EGFP expression in the germline of either male or female embryos up to stage 15 (Fig. S2B, S2F). To accommodate the possibility of a delay between the activation of the *SxlPE-*5.2kb transgene and our ability to detect EGFP expression (as is observed in the somatic cells), we also characterized its expression at later stages of development. However, we failed to observe any germline EGFP expression at the first, second, or third larval instar (L1, L2, or L3 respectively) stages (Fig. S2B-S2I). Thus, *SxlPE-*5.2kb also does not contain the *cis*-regulatory elements sufficient for *SxlPE* activity in the germline. Interestingly, *SxlPE*-5.2kb exhibited sex-specific EGFP expression generally in the soma until L1, and this expression persisted in the somatic gonad until L2 (Fig. S2F-S2H). This was unexpected as *SxlPE* has been reported to shut off in the soma following activation of dosage compensation in the early embryo. Our observations could be due to the stability of the EGFP reporter used or may point at a possible mechanism that keeps *SxlPE* active even after X chromosomes can no longer be counted.

We next generated a *SxlPE*-EGFP reporter that includes the 5.2kb sequence discussed above, as well as an additional 5kb downstream of *SxlPE* up to the start of Exon 4 (Fig. S2A) (*SxlPE*-10.2kb). Flies carrying this transgene showed sex-specific nuclear EGFP expression in the soma by stage 15 of embryogenesis (Fig. S2J-S2K). Excitingly, we also observed sex-specific nuclear EGFP expression in the female germline during the first larval instar (L1) stage (GFP immunostaining Fig. 2A-2B’, endogenous GFP and quantification, Fig. 6F,G,I). Together, these data suggest that *SxlPE* activity in the germline is sex-specific as it is in the soma, and that it requires additional *cis*-regulatory elements that lie downstream of *SxlPE*. Further, germline *SxlPE* activation appears to occur later in development than previously thought (L1), although it is possible that there is a delay between when *SxlPE* is actually activated and when we are first able to detect EGFP expression.

**Figure 2:**
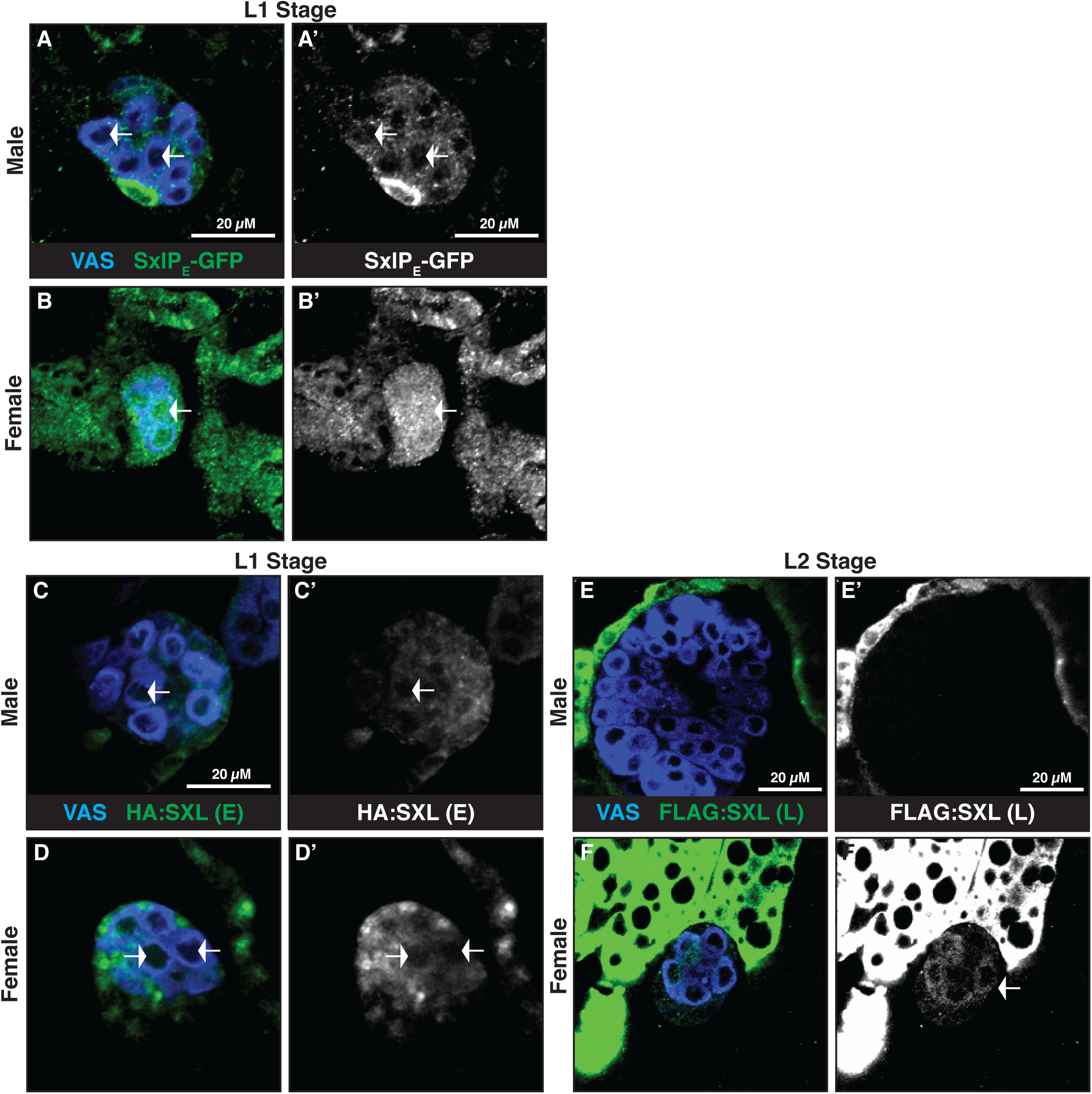
*SxlPE* is sex-specifically activated in the germline. A-B ’) Immunofluorescence of first instar (L1) gonads from flies bearing *SxlPE*-10.2kb reporter transgene. Note the presence of sex-specific nuclear GFP expression in female germ cells only. Arrows mark nuclei of germ cells. C-D’) Immunofluorescence of first instar (L1) gonads from flies bearing the HA:SxlE1 (Early Sxl) tag. Arrows mark cytoplasmic HA-background staining. E-F’) Immunofluorescence of second instar (L2) gonads from flies bearing the FLAG:SxlL2 (Late Sxl) tag. Arrows mark FLAG-positive germ cells. Note the presence of FLAG (Late Sxl) expression in female germ cells only. The anti-FLAG immunoreactivity in the fat body of both sexes is also present in wild-type stocks with no FLAG-tagged proteins, indicating it is antibody background. VAS stains germ cells.

### Late Sex lethal protein is expressed only after *SxlPE* is activated in the germline

Since *SxlPE* is sex-specifically activated in the germline, we hypothesized that, like in the soma, Early Sxl protein derived from *SxlPE* would be required to produce Late Sxl protein derived from *SxlPM*. To study expression of the Early and Late Sxl proteins in the germline, we used CRISPR-Cas9 genome editing to separately tag Sxl’s Early and Late protein products. We inserted a 2x human influenza hemagglutinin (2xHA) epitope at the N-terminus of the coding sequence of exon E1, which is specific to the mRNA produced by *SxlPE,* to generate a HA:SxlE1 ‘Early (E) Sxl’ tag (Fig. S3A). Separately, we inserted a 3xFLAG epitope at the N-terminus of the coding sequence of exon L2, which is specific to the mRNA produced by *SxlPM,* to generate a FLAG:SxlL2 ‘Late (L) Sxl’ tag (Fig. S3A) (Note: these tags are in separate stocks). Both of these tagged alleles of *Sxl* appear wildtype for *Sxl* function as homozygous females are viable and fertile.

Using anti-HA antibody, we were able to observe female-specific Sxl Early protein in somatic cells beginning at stage 9, and this expression was primarily nuclear, consistent with Sxl’s role in alternative splicing. Early Sxl expression in the soma persisted until the L1 stage (Fig. S3B-S3G), and no expression was observed after this time. In addition to the somatic gonadal tissue, we confirmed Early Sxl expression in the L1 stage in another somatic tissue, the developing gut (Fig. S3H-S3I). The continued presence of Early Sxl protein at the L1 stage is consistent with what we observed with the *SxlPE*-10.2kb reporter and suggests that Early Sxl protein is expressed, or at the very least maintained, much later in development than previously thought, and well after dosage compensation has been initiated. To our surprise, we were unable to detect anti-HA immunolabeling in the developing germline at any stage examined (embryo, and L1-L2). A low level of signal was observed in the cytoplasm of the germline (arrows in Fig. 2C-D’), but this was not clearly different between males and females, and may represent background staining. Detection of Early Sxl in the soma required tyramide amplification of the antibody signal, and so Early Sxl protein may be expressed at very low levels, which was undetectable in the germline even after tyramide amplification. Regardless, this prevents us from making a conclusion about Early Sxl protein expression in the germline.

Using the FLAG-tagged Late Sxl allele (FLAG:SxlL2), we observed anti-FLAG immunolabeling in female germ cells starting at the second larval instar (L2) stage (Fig. 2E-2F’). Female germ cells remained positive for anti-FLAG labeling through to adulthood and expression patterns were consistent with anti-Sxl antibody labeling (Fig. S3J-S3M’), which can be used to visualize Sxl in the germline starting at the third larval instar stage and onwards (Data not shown). No anti-FLAG immunoreactivity was observed in male germ cells. Though our RNA FISH analysis indicates that expression of *SxlPM* begins in the germline at stage 5, and is active in most germ cells by stage 10 (Fig. 1C), the *SxlPE* reporter (*SxlPE*-10.2kb) is only detectable in L1 germ cells (Fig. 2B), indicating that *SxlPE* expression begins much later in the germ cells than the soma. This is consistent with our ability to detect Late Sxl protein only in L2 germ cells; if Early Sxl protein is required to splice *SxlPM* transcripts to produce Late Sxl protein, then we would expect to detect Late Sxl protein only after *SxlPE* is active in the germline.

### *sisterless A* loss-of-function in the female germline results in ovarian tumors and germline loss

We next wanted to investigate the *trans-*acting factors that regulate *Sxl* in the germline. As discussed above, previous work indicates that germ cells use a different combination of XSEs than the soma. However, it remains possible that some of the individual somatic XSEs contribute to the X chromosome counting mechanism in the germline, perhaps along with unknown germline-specific XSEs. To investigate the potential role of the individual somatic XSEs in germline *Sxl* activation, we knocked down their expression specifically in the germline using RNAi (*nanos*-GAL4; UAS-*sisA* RNAi, UAS-*sisB* RNAi, UAS-*sisC* RNAi, or UAS-*runt* RNAi). We then immunolabeled adult gonads to look for phenotypes resembling *Sxl* loss of function, such as the formation of ovarian tumors. We found that RNAi knockdown of *sisB*, *sisC*, or *runt* in the germline did not result in any aberrant ovarian phenotypes (Fig. S4A-S4D’). Germ cells in these ovaries appear to differentiate properly as indicated by the progressive increase in the size of their nuclei, characteristic of polyploid nurse cells during oogenesis.

In contrast, *sisA* germline RNAi (*sisA* RNAi 1) resulted in ovaries that exhibited ovarian germ cell tumors (Fig. 3C-3C’, S5C, S5D) similar to those observed in *Sxl* LOF (Fig. 3B-3B’, S5C) or when XY germ cells develop in an XX soma [32,46–48]. While a high percentage of ovaries exhibited germline tumors (Fig. S5C), not all ovarioles in each ovary were tumorous. To confirm that this phenotype was not due to off-target effects sometimes seen with RNAi, we generated an additional UAS-*sisA* RNAi transgene (*sisA* RNAi 2) (Fig. S5D). Expression of this RNAi line in the germline also produced ovarian germ cell tumors (Fig. S5B-S5B’, S5D), but these tumors were less severe and occurred with lower frequency (Fig. S5C). An additional UAS-*sisA* RNAi transgene did not result in any observable phenotypes (Fig. S5D, Data not shown). When 2 copies of *nanos*-GAL4 were used to drive increased expression of UAS-*sisA* RNAi 1, we observed severe germline loss resulting in germ cell-less ovaries (Fig. 3D-3D’, S5C) and subsequent infertility. This is similar to what is observed in strong alleles of other sex determination genes such as *ovo* and *ovarian tumor (otu)*, as well as when XY germ cells develop in an XX soma [40,75–79].

**Figure 3:**
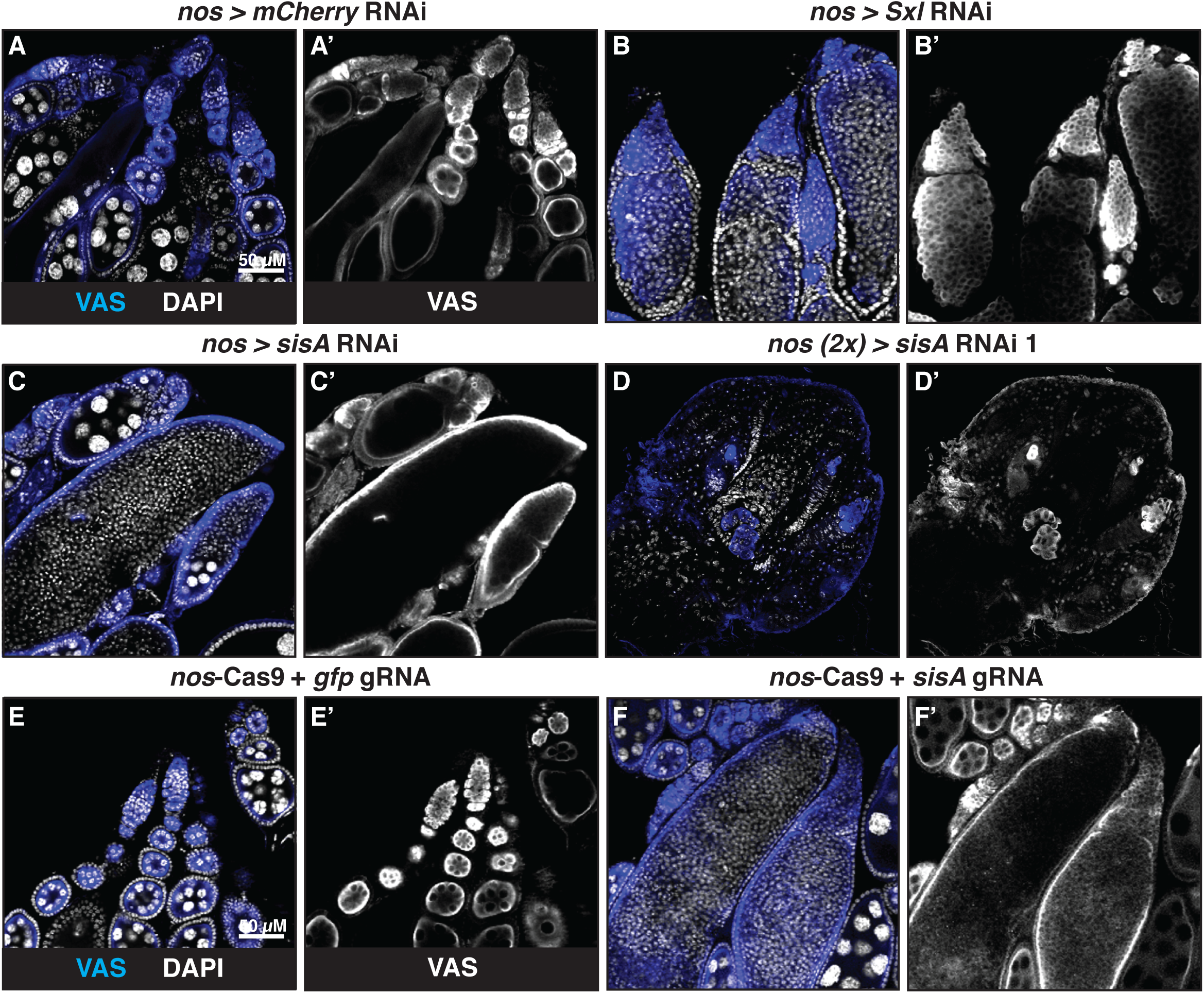
*sisA* loss of function in the female germline results in ovarian tumors and germ cell loss. A-F’) Immunofluorescence of adult ovaries to characterize germ cell phenotypes. A-A’) Wildtype ovary from fly with germline-specific *mCherry* RNAi as control. B-B’) Ovary with germ cell tumors from fly with germline-specific *Sxl* RNAi. Knockdown of *Sxl* causes a germline tumor phenotype. C-C’) Ovary with germ cell tumors from fly with germline-specific *sisA* RNAi (*sisA* RNAi 1). D-D’) Ovary with severe germ cell loss from fly with strong germline-specific *sisA* RNAi (2x GAL4). E-G) Animals expressing Cas9 in the germline along with guide RNAs for control (*gfp*) or *sisA* to create *de novo* “G0” mutations in the germline*. E*-E’) Wildtype ovary from fly with control guide RNAs for *gfp*. F-F’) Ovary with germ cell tumors from fly with guide RNAs generating mutations in *sisA*. G-G’) Germ cell-less ovary from fly with guide RNAs generating mutations in *sisA*. VAS stains germ cells. DAPI stains DNA (nucleus).

Next we wanted to investigate the null phenotype of *sisA* in the germline. Since homozygous *sisA* mutant females and hemizygous *sisA* mutant males are embryonic lethal [57], we generated *sisA* germline-specific loss of function mutations using tissue-specific CRISPR-Cas9 genome editing (“G0 CRISPR”, [69–71]). Male flies that ubiquitously express either 2 or 4 different guide RNAs (gRNAs) targeting the *sisA* gene region (Fig. S5D) were mated to females expressing Cas9 in the germline under the *nanos* promoter. F1 females from this cross also exhibited ovarian tumors as well as germ cell-less ovaries (Fig. 3E-3G’, S5C), similar to what was observed using RNAi (Fig. 3C-D’). Taken together, these data indicate that *sisA* is required for female germline differentiation and maintenance. Additionally, germline *sisA* loss of function is similar to *Sxl* loss of function, suggesting *sisA* is a candidate XSE for *SxlPE* activation in the germline.

### *sisterless A* expression in the embryonic germline precedes *Sex lethal* activation

In order for *sisA* to act as an XSE for activating *Sxl* in the germline, it should be expressed zygotically in germ cells prior to sex-specific activation of *SxlPE* in the germline. It has been reported that *sisA* mRNA is excluded from the nuclei that bud off to form pole cells [28]. However, reported microarray expression data demonstrates that *sisA* transcripts are enriched in PGCs at the 1-to-3 hour time point [80]. To observe zygotic transcription of *sisA* in the germline we utilized RNA FISH against nRNA being transcribed from the *sisA* locus. As a validation of our approach, we observed signals from *sisA* RNA FISH probes as fluorescent nuclear foci in somatic nuclei starting at nuclear cycle 8 (Fig. S6A-S6A’) and also at later stages in yolk cell nuclei, consistent with previous reports (Fig. S6B-S6B’) [28,57]. Interestingly, we were able to detect *sisA* RNA FISH signals in PGCs at stages 3-6 (3-5 shown in Fig. 4A-4C’). At later stages, the strong expression levels of *sisA* RNA in yolk cell nuclei made it difficult to distinguish fluorescent nuclear foci in neighboring PGCs. This suggests that *sisA* is indeed zygotically transcribed in the early embryonic germline. We quantified this expression in sexed embryos and found that *sisA* is expressed in the PGCs of both sexes (Fig. 4D).

**Figure 4:**
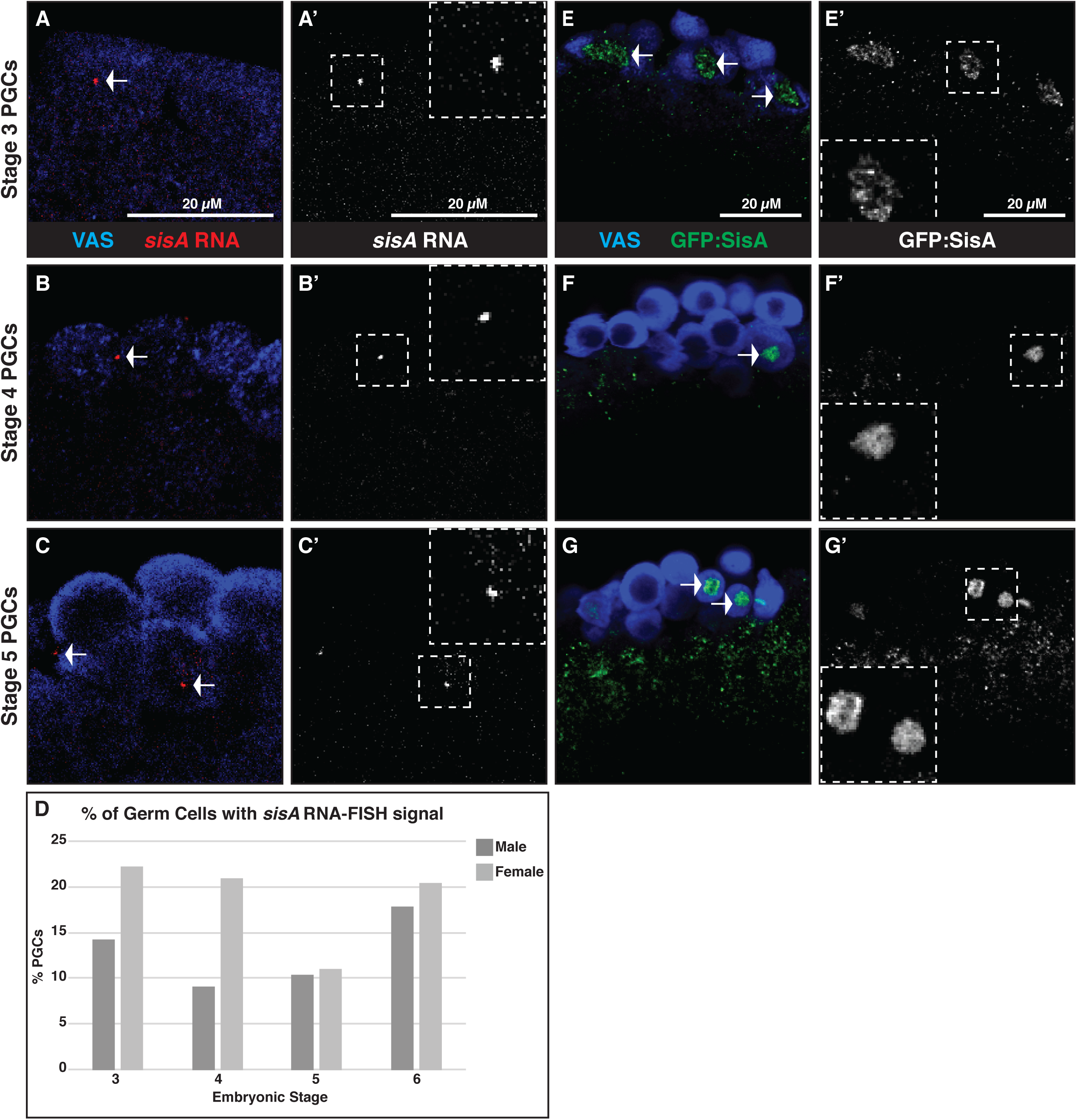
*sisA* is zygotically expressed in the germline. A-C ’) RNA-FISH (+ Immunofluorescence) against *sisA* in embryonic PGCs (Stages 3-5 shown). Smaller white dashed boxes specify the region zoomed in on, and presented in larger white dashed boxes. Arrows mark fluorescent nuclear foci of RNA-FISH signals. Note that a single focus is observed in female PGCs suggesting that X chromosomes may be paired in the germ cells. VAS stains germ cells. D) Graph showing percentage of PGCs with observable RNA-FISH probe signals between stages 3 and 11 of embryogenesis. *sisA* is zygotically expressed in PGCs of both sexes. E-G’) Immunofluorescence of embryonic PGCs (Stages 3-5 shown) from flies bearing sfGFP:SisA (SisA tag). Note that this expression is nuclear, consistent with SisA’s characterization as a bZIP transcription factor. Arrows mark GFP-positive germ cell nuclei. VAS stains germ cells.

We next wanted to observe SisA protein in the germline. Since no antibodies against SisA have been generated to date, we used CRISPR-Cas9 genome editing to insert a super-folder Green Fluorescent Protein (sfGFP) epitope tag at the N-terminus of the SisA protein (sfGFP:SisA) (Fig. S6C). Importantly, the epitope tag does not appear to affect SisA function as homozygous females and hemizygous males were viable and had no observable gonad phenotypes (Fig. S6D-S6E’). Furthermore, anti-GFP immunolabeling is restricted to the nucleus, which is consistent with the predicted function of SisA as a bZIP transcription factor. In somatic cells, we observed expression of sfGFP:SisA in pre-blastoderm syncytial nuclei (Fig. S6F-S6F’) and in yolk cell nuclei (Fig. S6G-S6G’), consistent with somatic *sisA* RNA expression.

Excitingly, similar to what we observed using RNA FISH, sfGFP:SisA expression was also observed in PGCs at stage 3, 4, and 5 (Fig. 4E-4G’). Additionally, this expression persisted in PGCs until stage 10 (Fig. S6H-S6H’). This late expression of SisA is consistent with its potential role in acting as an XSE for *SxlPE* activation in the germline. Like with *sisA* RNA, we were only able to detect sfGFP:SisA in a subset of PGCs in any one embryo. sfGFP:SisA expression was not observed later in first, second, or third larval instar stages, in either the germline or the soma (Data not shown). We conclude that SisA is expressed in the embryonic germline and this expression precedes the activation of *SxlPE*, which is consistent with a role as a germline XSE for the activation of *Sxl*.

### *sisterless A* is required for expression of Sxl in the female germline

If *sisA* acts as an activator of *Sxl* in the female germline, we would expect a loss of Sxl expression upon loss of *sisA* function. Strikingly, using anti-Sxl immunolabeling, we observed that the germ cell tumors in *sisA* RNAi ovaries had a strong reduction in Sxl expression when compared with controls (Fig. 5A-5C’, S7A-S7B’). Tumorous germ cells that were mutant for *sisA* using G0 CRISPR also had lower levels of Sxl expression (Fig. S7C-S7D’). Somatic Sxl expression remained unaffected in both cases. To examine Sxl expression specifically in those germ cells that express highest levels of Sxl (i.e. the early, undifferentiated germline), we performed *sisA* RNAi in the germline of *bag of marbles* (*bam*) mutants, which are enriched for germ cells robustly expressing Sxl (Fig 5D-5D’). We found that knocking down *sisA* using RNAi led to a dramatic reduction of Sxl antibody labeling in the germ cells of *bam* mutant ovaries (Fig. 5E-5E’), demonstrating that knocking down *sisA* leads to a loss of Sxl expression in the germline. To test whether *sisA* acts as a regulator of the Sxl PE promoter, we examined the effects of knocking down *sisA* function in the germline (*nos>sisARNAi)* on the expression of the Sxl PE-GFP reporter (*SxlPE*-10.2kb). Indeed, loss of *sisA* caused a significant reduction of germline GFP expression from the Sxl PE-GFP reporter, but did not reduce expression down to levels observed in males (Fig. 6F-I). This is consistent with SisA being one of, but not the only, regulator of Sxl PE expression in the germline.

**Figure 5:**
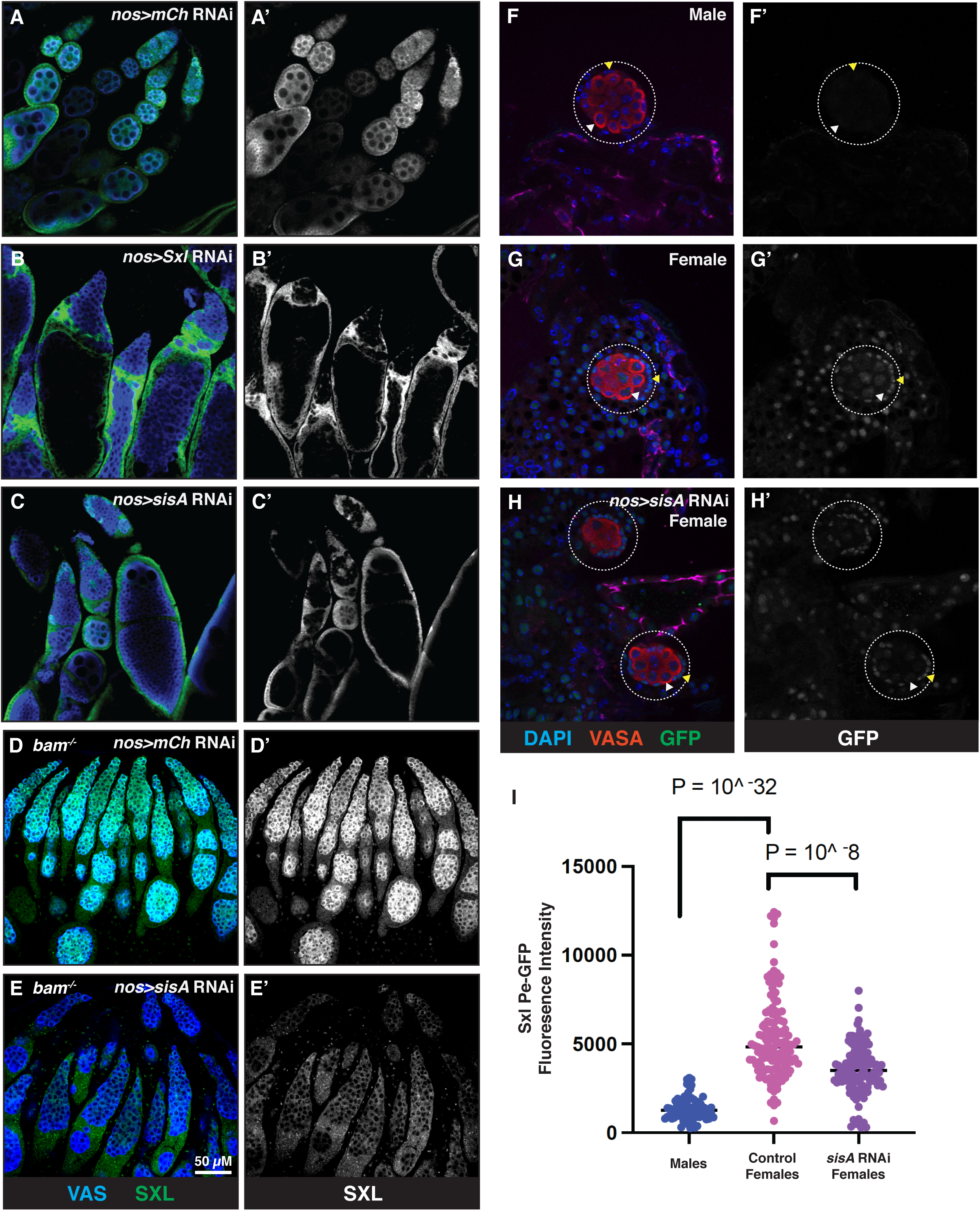
Sxl expression in the female germline depends on *sisA*. A-E’) Immunofluorescence of adult ovaries using anti-Sxl antibody (green). Anti-Vas stains the germ cells. A-A’) Wildtype ovary from fly with germline-specific *mCherry* RNAi. Sxl staining is highest in the early germ cells and decreases in differentiating germ cells. B-B’) Ovary with germ cell tumors from fly with germline-specific *Sxl* RNAi. Knockdown of *Sxl* causes a germline tumor phenotype and germ cells lack Sxl staining. C-C’) Ovary with germ cell tumors from fly with germline-specific *sisA* RNAi. Note that tumorous germ cells lack Sxl staining. Somatic Sxl remains unaffected. D-D’) *bam* mutant ovary with germline-specific *mCherry* RNAi. *bam* mutations cause a germline tumor phenotype and an expansion of germ cells that highly express Sxl. E-E’) *bam* mutant ovary with germline-specific *sisA* RNAi. Sxl staining is dramatically reduced in the germline. F-G’) Visualization of endogenous GFP expression (Green) from the *SxlPE*-10.2kb reporter in L1 stage males (F, F’), females (G, G’) and females expressing *sisA RNAi* in the germline (*nos>sisARNAi)*. White arrowheads indicate examples of germ cells and yellow arrowheads indicate somatic gonadal cells. H) Quantification of GFP fluorescence intensity in germ cells from samples as in F-G.

**Figure 6:**
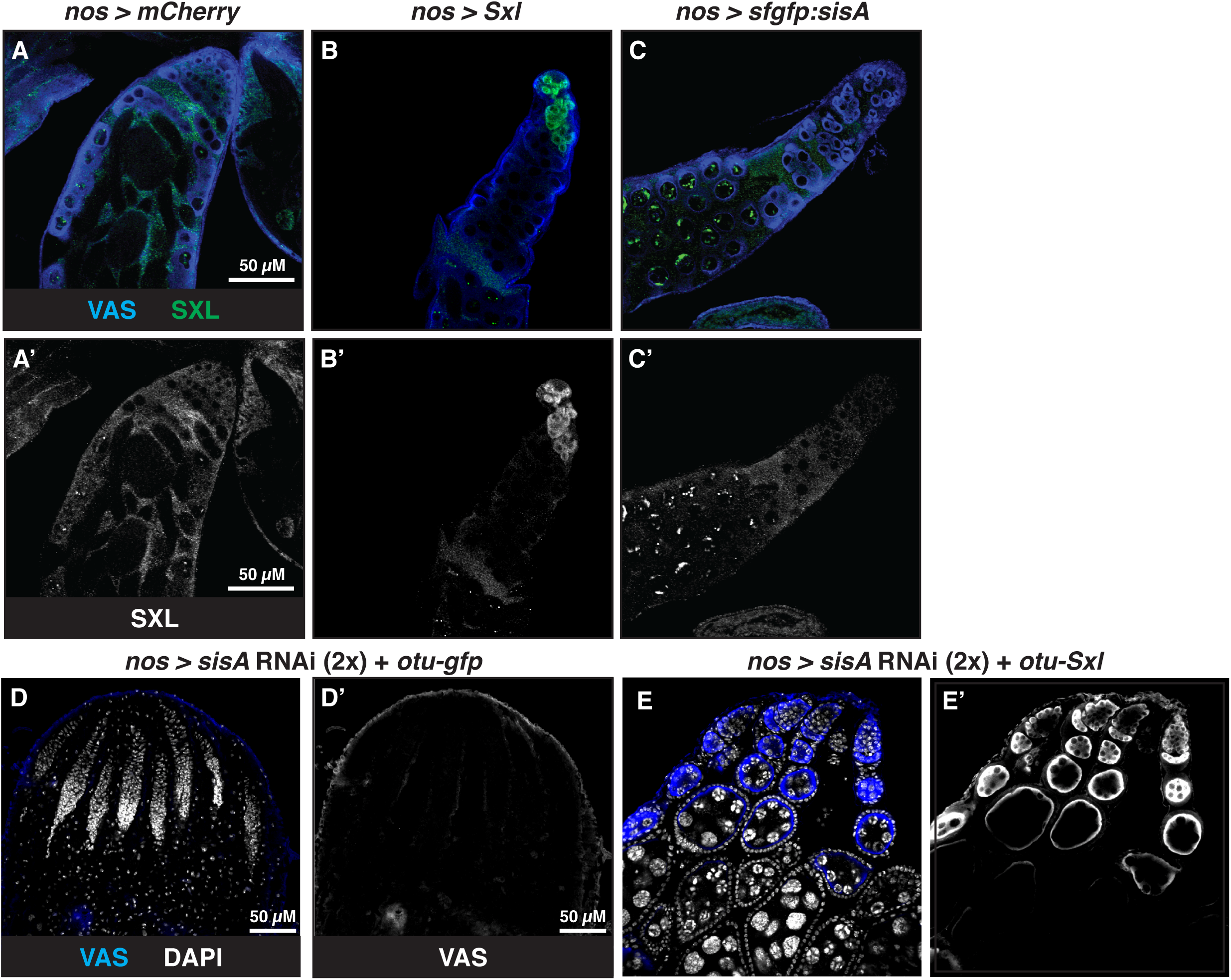
*sisA* is necessary but not sufficient to feminize the germline. A-C ’) Immunofluorescence of adult testes to characterize anti-Sxl staining. A-A’) Wildtype testis from fly with germline-specific overexpression of *gfp*. B-B’) Testis from fly with germline-specific overexpression of *Sxl*. Sxl staining is observed in the early germ cells. The anterior tip of the testis looks mildly atrophied. C-C’) Testis from fly with germline-specific overexpression of *sisA*. No Sxl staining is observed in the germ cells. The testis resembles the wildtype control. D-E) Immunofluorescence of adult ovaries to characterize rescue of *sisA* loss-of-function by *Sxl* expression. All animals strongly express UAS-*sisA* RNAi using two copies of *nos-*Gal4. D-D’) Germ cell-less ovary from *sisA-*RNAi flies with ectopic expression of a control protein (GFP) using the *otu* promoter. Flies of this genotype are sterile. E-E’) Ovary from *sisA-*RNAi flies with ectopic expression of *Sxl* under the control of the *otu* promoter. Note the rescue of the germline and the wildtype appearance of the ovary, indicating normal oogenesis. VAS stains germ cells. SXL stains Sxl. DAPI stains DNA (nucleus).

These data show that *sisA* is necessary for Sxl expression in the germline, and we next wanted to test whether ectopic expression of *sisA* would be sufficient to promote Sxl expression. Expression of UAS-*Sxl* in the male germline leads to inhibition of the JAK/STAT pathway and loss of germline stem cells ([86] and Fig. 6B-6B’). Expression of UAS-*sisA* did not lead to detectible expression of Sxl in the testis (Fig. 6C-6C’). This indicates that the overexpression of *sisA* alone is not sufficient to activate *Sxl* in the germline and that, like in the soma, additional XSEs are required to activate *SxlPE* in the germline.

If *sisA* is required for *Sxl* expression, and this is its primary role in the germline, then expression of *Sxl* should bypass the need for *sisA* in the germline and rescue the germline defects observed with *sisA* LOF. To test this we expressed *Sxl* under the control of a female germline-specific promoter (*otu*) in a *sisA* LOF background. We used the strongest *sisA* loss of function condition (2x *nos*-GAL4 > *sisA* RNAi 1) which results in severe germline loss, tumor formation, and infertility (Fig. 6D-6D’, S7E). Excitingly, 50% of *sisA* RNAi + *otu* > *Sxl* ovaries exhibited a wildtype germline morphology (Fig. 6E-6E’, S7E) and were fertile, suggesting *Sxl* was able to provide a robust rescue. Another 20% of these ovaries showed a partial rescue of the germline, but contained pervasive germline tumors (Fig. S7E). Taken together, these data indicate that *sisA* is necessary, but not sufficient, for *Sxl* expression in the female germline and that activation of *Sxl* is the primary germline role for *sisA*.

## DISCUSSION

### The sex-specific activation of *Sxl* in the germline

Previous evidence indicated that sex-specific activation of *SxlPE* is important for *Sxl* expression in germline, as it is in the soma. Transcripts from the *SxlPE* promoter were detected in early germ cells [50] and positive autoregulation of Sxl was observed in the germline [49], suggesting that the same mechanism of Early Sxl protein from *SxlPE* being required for splicing of *SxlPM* transcripts occurs in germ cells and soma. Our data also support this model. Using a *SxlPE* transcriptional reporter, we found that *SxlPE* is active in female germ cells by L1 (Fig. 2B-2B’). Further, while we observed activation the *SxlPM* promoter in embryonic germ cells (Fig. 1B-1B’’, 1C), we only observed expression of Late Sxl protein in the germ cells at L2 (Fig. 2F-2F’), after we observed activation of *SxlPE.* These data are consistent with *SxlPM* transcripts being unable to produce Sxl protein until after sex-specific activation of *SxlPE*. One difference between the germline and the soma appears to be the timing of *SxlPE* activation relative to that of *SxlPM*. In the soma, *SxlPE* is activated prior to *SxlPM*, while in the germline, there is no time at which we detect *SxlPE* alone by RNA FISH (Fig. 1C), and *SxlPE* is activated later in development, as judged by the *SxlPE* transcriptional reporter. This is still consistent with the autoregulatory model for *Sxl* activation, but suggests that the control of *SxlPE* activation is different in the germline than the soma.

We also found that the *cis-*regulatory elements that regulate female-specific *SxlPE* expression in the germ cells are different from those in the soma. Previously, it was reported that a 1.5kb genomic fragment just upstream of *SxlPE* was sufficient to regulate female-specific expression in both the soma [20,74] and the germline [50]. However, we were unable to observe germline expression from this *SxlPE* reporter or even a larger *SxlPE* reporter containing more of the upstream DNA (Fig. S2F-I). We only observed activation of *SxlPE* when we included an even larger genomic region, including sequences both upstream and downstream of *SxlPE* (Fig. S2A, 2B-2B’). These data do not exclude sequences within the 1.5 kb region upstream of *SxlPE* being important for its activation in the germline, but indicate that these sequences are not sufficient for activation, as they are in the soma. Similarly, sequences outside this 1.5 kb region may also augment expression in the soma. The fact that the *cis*-regulatory logic of *SxlPE* is different in the germline than the soma is consistent with previous work demonstrating that the combination of *trans-*acting factors, the XSEs, that activate *SxlPE* in the germline are different from the soma [33,52,81]. Future work will involve identifying the specific enhancers that regulate *SxlPE* expression in the germline, which could in turn help identify additional germline *trans*-regulators of *Sxl*. In summary, we have shown that the *cis-*regulatory logic of *Sxl* activation is different between the germline and the soma, however, the key step in determining the sex of both these cell types is the female-specific activation of *SxlPE*.

Finally, a surprising observation from our study was the presence of sex-specific *SxlPE* transcriptional reporter activity, as well as Early Sxl protein, in somatic cells as late as the L1 larval stage (Fig. S2F-S2G, S3E-S3F, S3I). In order for XSEs to act, they need to be expressed at higher levels in XX cells than XY cells. Once X chromosome dosage compensation is initiated at the late blastoderm stage, there should no longer be a difference in XSE expression between XX and XY cells. This is the entire logic behind the autoregulatory model for Sxl expression; *SxlPE* is activated by XSEs only in females, and is then turned off prior to the time when dosage compensation equalizes XSE expression. Why then is Early Sxl protein and *SxlPE* activity detected at the L1 stage? One possibility is that both the GFP reporter and the Early Sxl protein are stable and perdure to the L1 stage. An alternative possibility, however, is that some other mechanism, such as feedback regulation within the sex determination system, maintains *SxlPE* activity much longer than previously thought. Interestingly, *Sxl* contains a highly-conserved binding site for the Doublesex transcription factor which could facilitate such an XSE-independent feedback regulation of *SxlPE* [82].

### *sisterless A* as an activator of *Sxl*

While the combination of XSEs at work in the germline may be different than in the soma, our data also indicate that at least one XSE, SisA, is shared between them. Loss of *sisA* in the germline leads to the formation of germline tumors (and germline loss) (Fig. 3C-3D’, 3F-3G’, S5B-S5C) as well as a loss of Sxl expression in the female germline (Fig. 5C-5E’, S7B-S7D’). Additionally, we report that *sisA* is expressed in the germline prior to *SxlPE* (Fig. 4) and that the expression of Sxl is able to rescue the *sisA* loss-of-function phenotype in the germline (Fig. 6D-6E’, S7E), indicating that *sisA* lies upstream of *Sxl* in the germline sex determination cascade. Taken together, these data indicate that SisA is a germline activator of *Sxl* and may act as a germline XSE.

It remains to be seen whether SisA acts directly on *Sxl* in the germline or acts on other *Sxl* regulators, such as *ovo* or *otu*. Current thinking is that SisA is a transcriptional activator for *SxlPE* in the soma [9,28], and the simplest model would be that it acts similarly in the germline. However, the gap in timing we observe between *sisA* expression in the germline (Stage 3-10, Fig. 4E-4G’, S6H-S6H’) and our ability to detect *SxlPE* activation (L1 larvae, Fig. 2B-2B’) leaves open the possibility that there are intermediate steps between *sisA* and *SxlPE.* SisA is a somewhat unusual leucine zipper protein and its DNA binding activity and specificity have never been determined. Thus, there is no evidence that it regulates *SxlPE* directly in the soma or germline. The transient nature of SisA expression during embryogenesis makes it difficult to conduct chromatin immunoprecipitation (ChIP) experiments under native conditions. Instead, we expressed GFP:SisA in *Drosophila* S2 tissue culture cells for ChIP analysis, but this experiment did not identify SisA genomic binding sites. Lastly, yeast two-hybrid analysis has indicated that SisA might physically interact with the basic-helix-loop-helix protein Daughterless (Da) [83], but co-expressing GFP:SisA and Da in S2 cells also did not result in an identifiable ChIP signal. Thus, while our data clearly indicate that *sisA* is required for activation of *Sxl* in the germline, and that this is its primary role in the germline, evidence that SisA is a direct, transcriptional activator of *SxlPE* will require identification and study of a SisA-binding enhancer within the *Sxl* locus.

It is puzzling that *Sxl* is a critical regulator of both germline and somatic sex determination, and is regulated by a two-X dose in both cell types, yet its mechanism of activation and its role in these cell types is so different. It is somewhat simplifying that at least one factor, *sisA*, is important for activation of *Sxl* in both cell types. Yet evidence still suggests that other aspects of *Sxl* activation in these cells will be different. Further, the most significant Sxl targets in the soma, *transformer* for sexual identity and *msl-2* for dosage compensation, do not play a role in the germline [45,47]. Instead, distinct factors such as *Tdrd5l* and the Jak/Stat pathway, which controls *Phf7* expression, are regulated by *Sxl* in the germline [67,84-86]. It will be interesting to examine how these independent roles for *Sxl* came to be, and how conserved each role is in different species.

## MATERIALS AND METHODS

### Fly stocks

The following fly stocks were used: *SxlPE-EGFP* (BDSC# 24105), *SxlPE-EGFP* (BDSC# 32565), *vas*-Cas9 (BDSC# 51324), *vas*-Cas9 (BDSC# 56552), alphatub-piggyBac (BDSC# 32070), *nos*-Gal4 ([55]), *nos*-Cas9 (BDSC# 54591), *nos*-Cas9 (BDSC# 78782), UAS-*sisA*-RNAi 1 (TRiP.HMC03864, BDSC# 55181), UAS-*Sxl*-RNAi (TRiP.HMS00609, BDSC# 34393), UAS-*sisB*-RNAi (TRiP.GL01130, BDSC# 41594), UAS-*sisC*-RNAi (TRiP.HMS00545, BDSC# 33680), UAS-*run*-RNAi (TRiP.HMS01186, BDSC# 34707), UAS-*mCherry*-RNAi control (BDSC# 35785), *bam[1]* (Gift from A. Spradling, [56]), *bam[delta86]* (BDSC# 5427), *sisA[5]* (Gift from J. Erickson, [57]), UAS-*Sxl* (BDSC# 58484), *otu*-*GFP*.K10 (BDSC# 29727), *otu*-*GFP*.SV40 (BDSC# 29729), *otu*-Sxl (BDSC# 58491). Oregon R flies were used as wildtype flies, w[1118];Sco/CyO;MKRS/TM6B,Tb,Hu double balancers (Gift from X. Chen) were used for 2^nd^ and 3^rd^ chromosome balancing. FM7c was used for X chromosome balancing.

### Antibody staining and Tyramide Signal Amplification (TSA)

Adult testes were dissected in 1X PBS and fixed at room temperature for 20 minutes in 4.625% formaldehyde in PBS containing 0.1% Triton X-100 (PBTx). Adult ovaries, and larval gonads (first, second, and third instar) were dissected in 1X PBS and fixed at room temperature for 20 minutes in 5.25% formaldehyde in PBTx. Blocking and immunostaining was performed as previously described [58], and samples were mounted in 2.5% DABCO. Embryo collection, fixing, blocking and staining was performed as described previously [59]. All images were taken with a Zeiss LSM 700 confocal microscope. The following primary antibodies (sources) and concentrations were used: chicken anti-Vas 1:10,000 (K. Howard); rabbit anti-Vas 1:10,000 (R. Lehmann); mouse anti-Sxl 3:100 (M18, DSHB); mouse anti-Fas3 1:50 (7G10, DSHB); rat anti-NCad 3:100 (DN-Ex #8, DSHB); mouse anti-Lamin B 1:100 (ADL67.10, DSHB); mouse anti-Pros 1:10 (MR1A, DSHB); guinea pig anti-TJ 1:1,000 (J. Jemc); rabbit anti-GFP 1:1,000 (ab290, Abcam); mouse anti-FLAG 1:50 (F3165, Millipore Sigma); rat anti-HA 1:100 (ROAHAHA Clone 3F10, Roche); rabbit anti-HA 1:800 (C29F4 #3724, CST). DSHB: Developmental Studies Hybridoma Bank; CST: Cell Signaling Technologies. Secondary antibodies were used at 1:500 (Alexa Fluor, Host:Goat, Invitrogen).

TSA was performed using the manufacturer’s protocol (Tyramide SuperBoost Kits with Alexa Fluor Tyramides, Invitrogen). Larval gonads and embryos were fixed as previously described (See above). Samples were blocked in 10% Goat Serum for 1 hour at room temperature followed by incubation with a single primary antibody (target of signal amplification) as previously described [58,59] overnight at 4°C with nutation. Primary antibody was washed off using 1X PBS and samples were incubated with poly-HRP-conjugated secondary antibody for 1 hour at room temperature. Secondary antibody was washed off using 1X PBS and samples were incubated in tyramide working solution for 10 minutes at room temperature before stopping the reaction. Samples were washed with 1X PBS and immunolabeling with TSA was multiplexed with other antibodies following the standard immunolabeling protocol.

### Developmental Staging and Sexing

To obtain stage-specific embryos, embryos were pre-collected overnight on apple juice plates with fresh yeast paste and discarded to synchronize egg-laying. Embryos were then collected for 2 hours at 25°C after which the flies were removed and the embryos were aged as required. To obtain first (L1) and second (L2) instar larvae, embryos were aged 20 hours. The plates were cleared of any larvae that had hatched early. L1 larvae were collected after 4-8 hours. For L2 larvae, the L1 larvae were transferred to fresh plates with yeast paste and aged for an additional 24 hours. Third instar (L3) larvae were collected directly from vials. Sex of embryos was determined by counting the number of *Sex lethal* or *sisterless A* RNA FISH signals corresponding to number of X chromosomes or by somatic EGFP expression using the transcriptional *SxlP_E_-EGFP* reporters. Stage of the embryos was approximately determined by the position of the primordial germ cells (PGCs). Sex of the first and second larval instar gonads was determined by looking for presence or absence of a hub using anti-FasIII or anti-NCad immunolabeling. Sex of the third larval instar gonads was determined by morphology.

### Oligopaints Probe Design and Synthesis

Oligopaint probes for RNA FISH were designed using Oligopaints [60–62] by Dr. Kayla Viets from Dr. Robert Johnston’s laboratory, Department of Biology at Johns Hopkins University. Gene target sequences (*Sex lethal* and *sisterless A*) were run through an open-source bioinformatics pipeline made available by the Wu lab at Harvard Medical School (http://genetics.med.harvard.edu/oligopaints/) to identify sets of 50-bp (*Sex lethal*) or 30-bp (*sisterless A*) optimized probe sequences (libraries) to tile the nascent RNA transcript [63]. Library of Cy3-conjugated oligos against *sisterless A* RNA was ordered individually. Library of oligos against *Sex lethal* and its sub-libraries (*PM* vs. *PE+PM* probes) were ordered as part of a 90k oligopool including oligos for other gene targets (Gift from R. Johnston, Johns Hopkins University). To isolate target-specific probes (*Sex lethal*), five 19-bp barcording primers, target F and R; universal (univ) F and R; and sublibrary (sub) F were appended to the 5’ and 3’ ends of each probe.

PCR using target F and R primers allowed amplification of libraries of probes against target RNA from oligopool. PCR using sub F and target R primers allowed amplification of sub-libraries from whole target libraries. PCR using univ F and R primers allowed the conjugation of fluorophores (Cy3, Cy5), generation of single-stranded DNA (ssDNA) probes as well as addition of secondary sequences to allow amplification of RNA FISH signal using secondary probes (conjugated with fluorophores) that bind to primary probes. RNA FISH probes were generated as previously described [60–62]). Total number of probes per transcript are as follows: *sisA* RNA – 12; *SxlPM* RNA – 19; *SxlPE+PM* RNA – 36.

### RNA fluorescent in situ hybridization (FISH) and immunofluorescence

RNA FISH was performed using modified versions of the protocols described previously [60–63]. Oregon R embryos were collected (after a pre-collection was discarded) on apple juice plates with fresh yeast paste for 2 hours at 25°C and aged as needed. Embryos were collected, rinsed with 1X PBTx, and dechorionated using 50% bleach for 90 seconds and washed with 1X PBTx. Fixing was performed in scintillation vials in 50μL 10X PBS, 100μL 0.25M EGTA, 125μL fresh 16% formaldehyde, 225μL Milli-Q H_2_O, 1μL NP-40 (Tergitol solution), and 500μL heptane. Embryos were shaken vigorously by hand for 1 minute, then fixed for 20 minutes at room temperature with gentle agitation. The aqueous phase was removed and the embryos were devitellinized in by adding 500μL methanol and vigorous shaking for 2 minutes. Devitellinized embryos were collected washed three times in methanol. Embryos were rehydrated via 5-minute serial single washes in 75%, 50%, and 25% methanol in 1X PBTx. This was followed by three 5-minute washes in 1X PBTx and two 15-minute washes in 1X PBTx with 0.2U/μL RNase inhibitor. Embryos were blocked for 1 hour at room temperature in blocking buffer (1X PBTx + Western Blocking Reagent, 1:1) with nutation. They were then incubated in primary antibody diluted in 1X PBTx with 3% normal goat serum and 0.2U/μL RNase inhibitor overnight at 4°C with nutation. Primary antibody was rinsed off followed by three 20-minute washes with 1X PBTx. Embryos were then incubated with secondary antibody diluted in 1X PBTx with 3% normal goat serum and 0.2U/μL RNase inhibitor for two hours at room temperature. Secondary antibody was rinsed off followed by two 20-minute washes in 1X PBTx and one 20-minute wash in 1X PBS. Embryos were then washed as follows: four 5-minute washes in 2X SSCT, one 10-minute wash in 20% formamide in 2X SSCT, one 10-minute wash in 50% formamide in 2X SSCT, one 4-hour wash in 50% formamide in 2X SSCT at 37°C with shaking. Embryos were incubated with primary probe at a concentration of ≥5 pmol fluorophore/μL in hybridization buffer (50% formamide in 2X SSCT with 10% dextran sulfate (w/v) + 0.2U/μL RNase inhibitor) for 16-20 hours at 37°C with shaking. Primary probe was rinsed and followed by a 1-hour wash in 50% formamide in 2X SSCT at 37°C with shaking. Secondary probes were hybridized for 1 hour at 37°C with shaking at a concentration of ≥5 pmol fluorophore/μL in 50% formamide in 2X SSCT and 0.2U/μL RNase inhibitor. This was followed by two 30-minute washes in 50% formamide in 2X SSCT at 37°C, one 10-minute wash in 20% formamide in 2X SSCT, two 10-minute washes in 2X SSCT at room temperature, and one 10-minute wash in 2X SSC at room temperature. Embryos were washed for 10 minutes in 1X PBTx containing DAPI and incubated at room temperature for 10 minutes in SlowFade Diamond with DAPI, followed by mounting. All images were taken with a Zeiss LSM 700 confocal microscope.

### SxlPE reporter transgenes and constructs

*SxlPE*-10.2kb, a 10.2kb genomic sequence from a *Sxl* genomic clone BAC# CH321-74P19 (BACPAC Resources Center) cloned into the pJR16 vector in two steps (Gift from R. Johnston, Johns Hopkins University) using HiFi DNA Assembly (New England Biolabs). pJR16 uses an EGFP reporter with a nuclear localization sequence (nls). Fragment 1 extended from 116 bp downstream of *SxlPE* TSS to 5,116 bp upstream of *SxlPE* TSS and was assembled into pJR16 digested with AgeI and AscI to generate *SxlPE*-5.2kb. Fragment 2 extended from 117 bp downstream of *SxlPE* TSS to 5,108 bp downstream of *SxlPE* TSS and was assembled into Fragment 1 + pJR16 digested with AgeI.

Fragment 1 *SxlPE*-10.2kb Fw (5’-3’) – CCACCCCGGTGAACAGCTCCTCGCCCTTGCTCACCATGGTGGCGACCGGTAATGGGATAATCACAAAGTT
Fragment 1 *SxlPE*-10.2kb Rv (5’-3’) – CATGCTGCAGCAGATCTGGTCTAGAGCCCGGGCGAATTCGCCGGCGCGCCGTAATTTTTCTTTGCTCCTCCTG
Fragment 2 *SxlPE*-10.2kb Fw (5’-3’) – CAGCTCCTCGCCCTTGCTCACCATGCTGTACGATGAATCGA
Fragment 2 *SxlPE*-10.2kb Rv (5’-3’) – GAAAAACGTAACTTTGTGATTATCCCATTATGGATTTCAATTTTGATAC

The primers ensured that EGFP remained in frame with Exon E1’s coding sequence (CDS). Constructs were injected into embryos and integrated via PhiC31 integrase-mediated transgenesis (done at BestGene Inc.) into the same genomic location at P{CaryP}attP40 on Chromosome II.

### Generation of sisA RNAi 2 and 3

Transcript-specific knockdown was achieved using the VALIUM vector system (PMID: 21460824). A 21-nucleotide (nt) sequence without any off-targets greater than 16nt was selected based on the algorithm of [64]. The following sequences were selected:

sisA-RNAi 2 Sense – CGCCGACGAGGAGCAACGCUA
sisA-RNAi 2 Antisense – UAGCGUUGCUCCUCGUCGGCG
sisA-RNAi 3 Sense – CCGGUUCUGGUUCGGAUGUCA
sisA-RNAi 3 Antisense – UGACAUCCGAACCAGAACCGG

The top and bottom strands for the short hairpin were as follows:

sisA-RNAi 2 Top Strand – CTAGCAGTCGCCGACGAGGAGCAACGCTATAGTTATATTCAAGCATATAGCGTTGCTCCTCGTCGGCGGCG
sisA-RNAi 2 Bottom Strand – AATTCGCCGCCGACGAGGAGCAACGCTATATGCTTGAATATAACTATAGCGTTGCTCCTCGTCGGCGACTG
sisA-RNAi 3 Top Strand – CTAGCAGTCCGGTTCTGGTTCGGATGTCATAGTTATATTCAAGCATA-TGACATCCGAACCAGAACCGGGCG
sisA-RNAi 3 Bottom Strand – AATTCGCCCGGTTCTGGTTCGGATGTCATATGCTTGAATATAACTATGACATCCGAACCAGAACCGGACTG

The top and bottom strands were annealed and cloned into the VALIUM20 vector #1467 (Drosophila Genomics Resource Center, DGRC). Constructs were injected into embryos and integrated via PhiC31 integrase-mediated transgenesis (done at BestGene Inc.) into the same genomic location at P{CaryP}attP40 on Chromosome II. For RNAi-mediated germline-specific knockdown of *sisA*, *sisB, sisC, runt, Sxl, Ovo,* and *mCherry*, male flies carrying shRNA transgenes were mated with *nanos*-GAL4:VP16 virgin females [55] and the crosses were maintained at 29°C. The progeny were reared at 29°C until 3-5 days post-eclosion (unless otherwise specified).

### CRISPR-tagging

sfGFP:sisA, 2xHA:SxlE1, and 3xFLAG:SxlL2 were generated using scarless genome editing described in [65] to generate N-terminal tags. Guide RNA (gRNA) sequences were selected using the FlyCRISPR algorithm (http://flycrispr.molbio.wisc.edu) [66], contain 20 nucleotides each and have no predicted off-targets. The following gRNAs were selected:

sfGFP:sisA gRNA Target (5’-3’) – GTCCAATGGCAAGCTACCTG
2xHA:SxlE1 gRNA Target (5’-3’) – GCCTCCTTCGATCTTCTACC
3xFLAG:SxlL2 gRNA Target (5’-3’) – GACTTGTTGTTGTAGCCATA

The guides were cloned into the pU6-2-BbsI-gRNA vector #1363 (DGRC) using the listed primers that were phosphorylated using T4 polynucleotide kinase (PNK).

sfGFP:sisA gRNA Target sense (5’-3’) – CTTCGTCGTTGGCCAATCCGGATGC
sfGFP:sisA gRNA Target antisense (5’-3’) – AAACGCATCCGGATTGGCCAACGAC
2xHA:SxlE1 gRNA Target sense (5’-3’) – CTTCGCCTCCTTCGATCTTCTACC
2xHA:SxlE1 gRNA Target antisense (5’-3’) – AAACGGTAGAAGATCGAAGGAGGC
3xFLAG:SxlL2 gRNA Target sense (5’-3’) – CTTCGACTTGTTGTTGTAGCCATA
3xFLAG:SxlL2 gRNA Target antisense (5’-3’) – AAACTATGGCTACAACAACAAGTC

Donor plasmids to facilitate homology dependent repair were generated. The coding sequences of the different tags are cloned from genomic DNA of *vas*-Cas9 expressing flies (BDSC# 51324, 56552) adjacent to a piggyBac transposon that contained a DsRed expression construct resulting in a selectable-tagging-cassette in vectors pHD-sfGFP-ScarlessDsRed #1365, pHD-2xHA-ScarlessDsRed #1366, pHD-3xFLAG-ScarlessDsRed #1367 (DGRC). Approximately 1kb 5’ and 3’ Homology arms were assembled using HiFi DNA Assembly (New England Biolabs) upstream and downstream of this cassette with the following primers (5’-3’):

sfGFP:sisA 5’ Arm Fw (5’-3’) – GAATTCGCCAAAGGGATTTC
sfGFP:sisA 5’ Arm Rv (5’-3’) – CCGGAACCTCCAGATCCACCGGTGATTTTTTTCGATGTGTG
sfGFP:sisA cassette Fw (5’-3’) – ACACATCGAAAAAAATCACCGGTGGATCTGGAGGTTCC
sfGFP:sisA cassette Rv (5’-3’) – AAGTAAAGATGACTCCGTTCGGAACCTCCTGAACCACC
sfGFP:sisA 3’ Arm Fw (5’-3’) – CTGGTGGTTCAGGAGGTTCCGAACGGAGTCATCTTTACTTGCC
sfGFP:sisA 3’ Arm Rv (5’-3’) – GGTACCGCATTGGCCCAATTC
2xHA:SxlE1 5’ Arm Fw (5’-3’) – GCAATCTGTGTTCTTGGTATTTTG
2xHA:SxlE1 5’ Arm Rv (5’-3’) – GGAACATCGTATGGGTACATAATGGGATAATCACAAAGTTAC
2xHA:SxlE1 cassette Fw (5’-3’) – AACTTTGTGATTATCCCATTATGTACCCATACGATGTTCC
2xHA:SxlE1 cassette Rv (5’-3’) – ACAGTATCAAAATTGAAATCGGAACCTCCTGAACCACC
2xHA:SxlE1 3’ Arm Fw (5’-3’) – CTGGTGGTTCAGGAGGTTCCGATTTCAATTTTGATACTGTGAC
2xHA:SxlE1 3’ Arm Rv (5’-3’) – CGATCGAAGGTGAGTTTC
3xFLAG:SxlL2 5’ Arm Fw (5’-3’) – CGACCATGTCGTCCTACTATAAC
3xFLAG:SxlL2 5’ Arm Rv (5’-3’) – TCATGGTCTTTGTAGTCCATATCCTGAGAGTTGGGAGTG
3xFLAG:SxlL2 cassette Fw (5’-3’) – ACACTCCCAACTCTCAGGATATGGACTACAAAGACCATGAC
3xFLAG:SxlL2 cassette Rv (5’-3’) – CCCGGATTATTGTTGCCGTAGGAACCTCCTGAACCACC
3xFLAG:SxlL2 3’ Arm Fw (5’-3’) – CTGGTGGTTCAGGAGGTTCCTACGGCAACAATAATCCG
3xFLAG:SxlL2 3’ Arm Rv (5’-3’) – GCTAATGAGGGGATTCCTATG

Assembled donor constructs were cloned into pCR2.1-TOPO (TOPO TA Cloning Kit, ThermoFisher). Site-specific mutagenesis was also used to mutate the PAM sites in each donor plasmid. For sfGFP:sisA, the included linker sequence between 3’ UTR and the start of sfGFP was removed using site-specific mutagenesis (QuikChange, Agilent). The gRNA expression plasmid and donor plasmid were injected by BestGene into embryos of Vas-Cas9 flies (#56552, #51324, BDSC). DsRed positive flies were then crossed to a piggyBac transposase expressing line (#32070, #32073, BDSC) to excise DsRed resulting in an in-frame fusion of the tags’ and the respective genes’ coding sequences. Successful generation of CRISPR-tags was confirmed by sequencing.

### Generation of overexpression constructs

UAS-*sfGFP*, UAS-*sisA*, and UAS-*da* were generated by cloning the ORFs of *sfGFP*, *sisA*, and *da-PD* into pUASpB [67] using the listed primers. pUASpB is a modified version of pUASP [68] including an attB site for phiC31-mediated integration. UAS-*sfGFP:sisA* was generated by cloning a flexible linker sequence followed by the ORF of *sisA* (lacking a start codon) using the listed primers into UAS-*sfGFP* that had been linearized with SpeI. Constructs were assembled using HiFi DNA Assembly (New England Biolabs).

UAS-*sfGFP* Fw (5’-3’) – TACCCGCCCGGGGATCAGATCCGCGGCCGCATGGTGTCCAAGGGCGAG
UAS-*sfGFP* Rv (5’-3’) – GACTCTAGAGGATCCAGATCCACTAGTTCACTTGTACAGCTCATCCATGC
UAS-*sfGFP:sisA* Fw (5’-3’) –GCCGGCATCACCCTGGGCATGGATGAGCTGTACAAGATTAAGGCCGGCGGGT CG
UAS-*sfGFP:sisA* Rv (5’-3’) – ACGTTAACGTTCGAGGTCGACTCTAGAGGATCCAGATCCATCACTGCTCCATTTCCAGGC
UAS-*sisA* Fw (5’-3’) – CTGTTCATTGGTACCCGCCCGGGGATCAGATCCGCATGGAACGGAGTCATCTTTACTTGC
UAS-*sisA* Rv (5’-3’) – ACGTTAACGTTCGAGGTCGACTCTAGAGGATCCAGATCCATCACTGCTCCATTTCCAGGC
UAS-*da-PD* Fw (5’-3’) – AGGTCCTGTTCATTGGTACCCGCCCGGGGATCAGATCCGCATGGCGACCAGTGACGATG
UAS-*da-PD* Rv (5’-3’) – ACGTTAACGTTCGAGGTCGACTCTAGAGGATCCAGATCCATTAAAAGTGTTGTACATTTTGTAGGGG

Constructs flies were injected into embryos and integrated via PhiC31 integrase-mediated transgenesis (done at BestGene Inc.) into the same genomic location at P{CaryP}attP40 on Chromosome II. For germline-specific overexpression, male flies carrying UAS transgenes were mated with *nanos*-GAL4:VP16 virgin females [55] and the crosses were maintained at 29°C. The progeny were reared at 29°C until 3-5 days post-eclosion (unless otherwise specified).

### G0 CRISPR (Tissue-specific CRISPR)

Guide RNA (gRNA) sequences were selected using the FlyCRISPR algorithm (http://flycrispr.molbio.wisc.edu) [66], contain 20 nucleotides each and have no predicted off-targets. For *sisA*, four different gRNA sequences were selected, one within the coding region (Target 4) and one within the 3’ UTR (Target 3), and two upstream of the gene locus (Targets 1 and 2). As a control, four different gRNA sequences were selected against GFP. The following gRNAs were selected:

sisA-G0 gRNA Target 1 (5’-3’) – TCGTTGGCCAATCCGGATGCAGG
sisA-G0 gRNA Target 2 (5’-3’) – CACTGAGTCTACCTGATAATTGG
sisA-G0 gRNA Target 3 (5’-3’) – ATAGTGTAGCTATGTGTCGCAGG
sisA-G0 gRNA Target 4 (5’-3’) – GCTGAAAACGGAGCTTGCTATGG
gfp-G0 gRNA Target 1 (5’-3’) – CAGGGTCAGCTTGCCGTAGG
gfp-G0 gRNA Target 2 (5’-3’) – AGCACTGCACGCCGTAGGTC
gfp-G0 gRNA Target 3 (5’-3’) – CGGCCATGATATAGACGTTG
gfp-G0 gRNA Target 4 (5’-3’) – CATGCCGAGAGTGATCCCGG

Four guides (or two) were cloned into the pCFD5 vector #73914 (Addgene) as previously described [69] using the listed primers.

sisA-G0-4-PCR1-Fw (5’-3’) – GCGGCCCGGGTTCGATTCCCGGCCGATGCATCGTTGGCCAATCCGGATGCGTTTTAGAGCTAGAAATAGCAAG
sisA-G0-4-PCR1-Rv (5’-3’) – ATTATCAGGTAGACTCAGTGTGCACCAGCCGGGAATCGAACCC
sisA-G0-4-PCR2-Fw (5’-3’) – CACTGAGTCTACCTGATAATGTTTTAGAGCTAGAAATAGCAAG
sisA-G0-4-PCR2-Rv (5’-3’) – GCGACACATAGCTACACTATTGCACCAGCCGGGAATCGAACCC
sisA-G0-4-PCR3-Fw (5’-3’) – ATAGTGTAGCTATGTGTCGCGTTTTAGAGCTAGAAATAGCAAG
sisA-G0-4-PCR3-Rv (5’-3’) –ATTTTAACTTGCTATTTCTAGCTCTAAAACTAGCAAGCTCCGTTTTCAGCTGCACCAGCCGGGAATCGAACCC
GFP-G0-4-PCR1-Fw (5’-3’) – GCGGCCCGGGTTCGATTCCCGGCCGATGCACAGGGTCAGCTTGCCGTAGGGTTTTAGAGCTAGAAATAGCAAG
GFP-G0-4-PCR1-Rv (5’-3’) – GACCTACGGCGTGCAGTGCTTGCACCAGCCGGGAATCGAACCC
GFP-G0-4-PCR2-Fw (5’-3’) – AGCACTGCACGCCGTAGGTCGTTTTAGAGCTAGAAATAGCAAG
GFP-G0-4-PCR2-Rv (5’-3’) – CAACGTCTATATCATGGCCGTGCACCAGCCGGGAATCGAACCC
GFP-G0-4-PCR3-Fw (5’-3’) – CGGCCATGATATAGACGTTGGTTTTAGAGCTAGAAATAGCAAG
GFP-G0-4-PCR3-Rv (5’-3’) –ATTTTAACTTGCTATTTCTAGCTCTAAAACCCGGGATCACTCTCGGCATGTGCACCAGCCGGGAATCGAACCC

Constructs were injected into embryos and integrated via PhiC31 integrase-mediated transgenesis (done at BestGene Inc.) into the same genomic location at P{CaryP}attP40 on Chromosome II. For germline-specific CRISPR-mediated knockout of *sisA* and *gfp*, male flies carrying gRNA expressing transgenes were mated with *nanos*-Cas9 virgin females ([70], BDSC# 54591 or [71], BDSC# 78782) and the crosses were maintained at 29°C. The progeny were reared at 29°C until 3-5 days post-eclosion (unless otherwise specified).

### Generating sisA mutant

CRISPR-Cas9 was used as previously described to generate a *sisA* deletion mutant [72,73]. 2 guide RNA (gRNA) sequences were selected flanking the *sisA* gene using the FlyCRISPR algorithm (http://flycrispr.molbio.wisc.edu) [66], contain 20 nucleotides each, and have no predicted off-targets. The following gRNAs were selected:

sisA-del gRNA Target 1 (5’-3’) – GTCCAATGGCAAGCTACCTG
sisA-del gRNA Target 2 (5’-3’) – GATTACCTTCGGCCAGGCGA
The guides were cloned into the pU6-2-BbsI-gRNA vector #1363 (DGRC) using the listed primers that were phosphorylated using T4 PNK.
sisA-del gRNA Target 1 sense (5’-3’) – CTTCGTCGTTGGCCAATCCGGATGC
sisA-del gRNA Target 1 antisense (5’-3’) – AAACGCATCCGGATTGGCCAACGAC
sisA-del gRNA Target 2 sense (5’-3’) – CTTCGCACTGAGTCTACCTGATAAT
sisA-del gRNA Target 2 antisense (5’-3’) – AAACATTATCAGGTAGACTCAGTGC

A donor plasmid to facilitate homology dependent repair was generated. A 5’ homology arm and a 3’ homology arm were PCR-amplified from genomic DNA of *vas*-Cas9 expressing flies (BDSC# 51324, 56552) and cloned into multiple cloning sites found upstream and downstream of a removable 3xP3-DsRed marker in the pHD-DsRed-attP vector #1361 (DGRC).

sisA-del 5’ Arm Fw (5’-3’) – AGCACACCTGCACGACCGATGAAAATGGAGCAAGTGGAAAGCACACCTGCACGACCGATGAAAATGGAGCAAGTGGAA
sisA-del 5’ Arm Rv (5’-3’) – AGCACACCTGCACGATTAACCTTAGGCAATATGTCAGCCAGCACACCTGCAC GA TTAACCTTAGGCAATATGTCAGCC
sisA-del 3’ Arm Fw (5’-3’) – GGCGGCTCTTCCTAAAAAACGAATGCTTTCTTATTGGCGGCTCTTCCTAA AAAACGAATGCTTTCTTATT
sisA-del 3’ Arm Rv (5’-3’) – GGCGGCTCTTCCCGGTTTAATATACATATATTTGTGGCGGCTCTTCCCGG TTTAATATACATATATTTGT

The gRNA expression plasmid and donor plasmid were injected by BestGene into embryos of Vas-Cas9 flies (#56552, #51324, BDSC). DsRed-positive flies were balanced without DsRed removal to maintain a selectable marker. Successful deletion of the *sisA* gene and replacement with the DsRed marker was confirmed by sequencing.

## AUTHOR CONTRIBUTIONS

Raghav Goyal: Conceptualization, Investigation, Formal Analysis, Writing (original draft and editing), Funding Acquisition. Ellen Baxter: Investigation, Formal Analysis, Project Administration. Mark Van Doren: Conceptualization, Writing (review and editing), Supervision, Project Administration, Funding Acquisition.

## ACKNOWLEDGEMENTS

We thank VDRC, Vienna, and the Bloomington *Drosophila* Stock Center, Indiana, for flies; the Developmental Studies Hybridoma Bank for antibodies; and Flybase (www.flybase.org) for essential information. We thank Integrated Imaging Center at The Johns Hopkins University, Baltimore, for imaging. This work was supported by NSF Fellowship DGE-123285 (to R.G.) NIH grant R01GM113001 (to M.V.D).

## FIGURE LEGENDS

**Figure S1:**
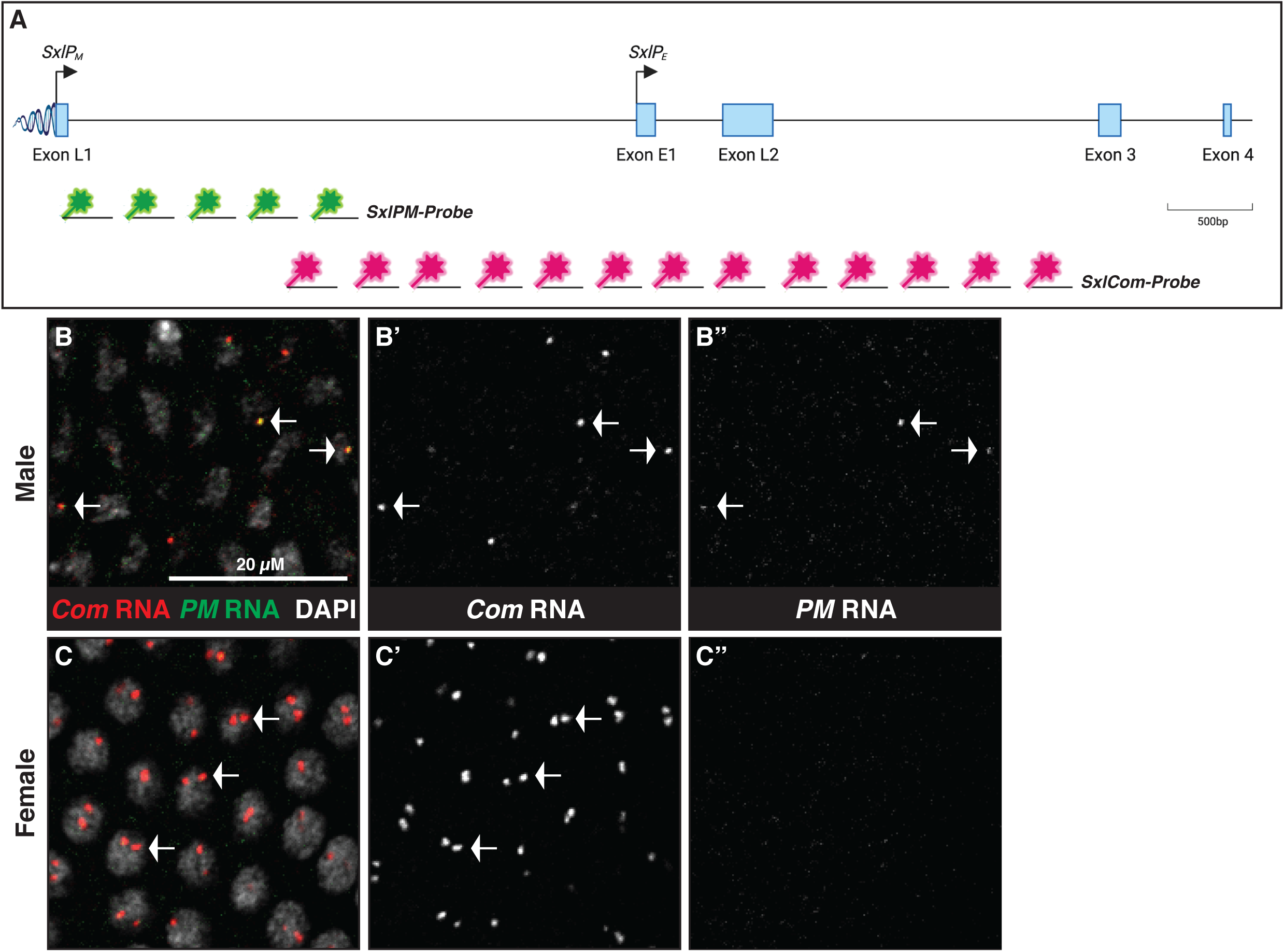
RNA-FISH against nascent RNA can be used to study *Sxl* promoter initiation. A) Cartoon showing relevant portion of the *Sxl* gene locus and span of *SxlPM* and *SxlPE+SxlPM* transcripts probed by *PM*-probes (Cy5, false colored Green) and *PE+PM*-probes (Cy3, Red) respectively, using RNA-FISH. Locus is drawn to scale. Size of probes is not drawn to scale. CDS: Coding sequences. UTR: Untranslated Region. B-C’’) RNA-FISH (+ Immunofluorescence) against the transcript from *SxlPM* only (*PM* RNA) versus the common transcript from *SxlPE* and *SxlPM* (*PE+PM* RNA) in embryonic stage 3 somatic cells. B-B’’) Male somatic cells always show both *PE+PM* and *PM*-probe signals. Note that a single focus is observed indicating a single X chromosome. C-C’’) Female somatic cells first have *PE+PM*-probe signals, indicative of *SxlPE* activity alone. Note that two foci are observed per nucleus indicating the presence of two X chromosomes that are unpaired. Arrows mark fluorescent nuclear foci of RNA-FISH signals except in C’’ which has no RNA-FISH signals. *PE+PM* RNA is probed by *PE+PM*-probes. *PM* RNA is probed by *PM*-probes. DAPI stains DNA (nucleus).

**Figure S2:**
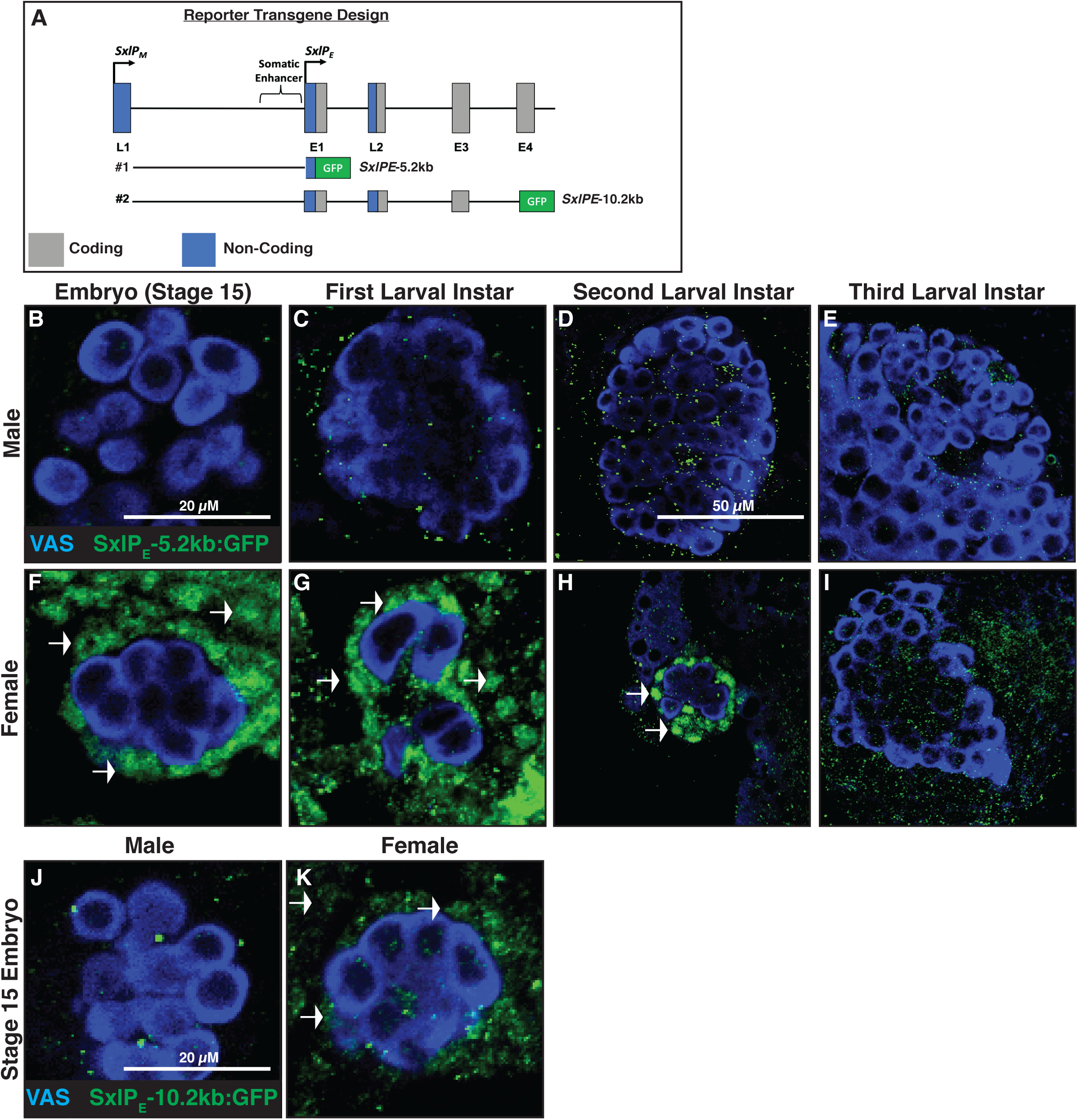
*SxlPE* requires different *cis*-regulatory elements in the soma and the germline. A) Cartoon showing relevant portion of the *Sxl* gene locus and design of transcriptional reporter constructs specific for *SxlPE*. EGFP reporters include a nuclear localization sequence (nls) to aid in visualization of expression. The 1.5kb somatic enhancer is illustrated. In *SxlPE*-5.2kb, EGFPnls replaces the CDS of Exon E1. In *SxlPE*-10.2kb, EGFPnls replaces the CDS of Exon 4. Locus is drawn to scale. EGFPnls is not drawn to scale. CDS: Coding sequences. UTR: Untranslated Region. B-I) Immunofluorescence of developing gonads to characterize *SxlPE-*5.2kb expression from stage 15 of embryogenesis to the third larval instar stage (L3). Note the presence of sex-specific nuclear GFP expression in female somatic cells. No GFP expression is observed in the germ cells at any stage. Arrows mark GFP-positive somatic cells. J-K) Immunofluorescence of embryonic stage 15 gonads to characterize *SxlPE*-10.2kb expression. Note the absence of GFP expression in female germ cells but presence of GFP expression in female somatic cells. Arrows mark GFP-positive somatic cells. VAS stains germ cells.

**Figure S3:**
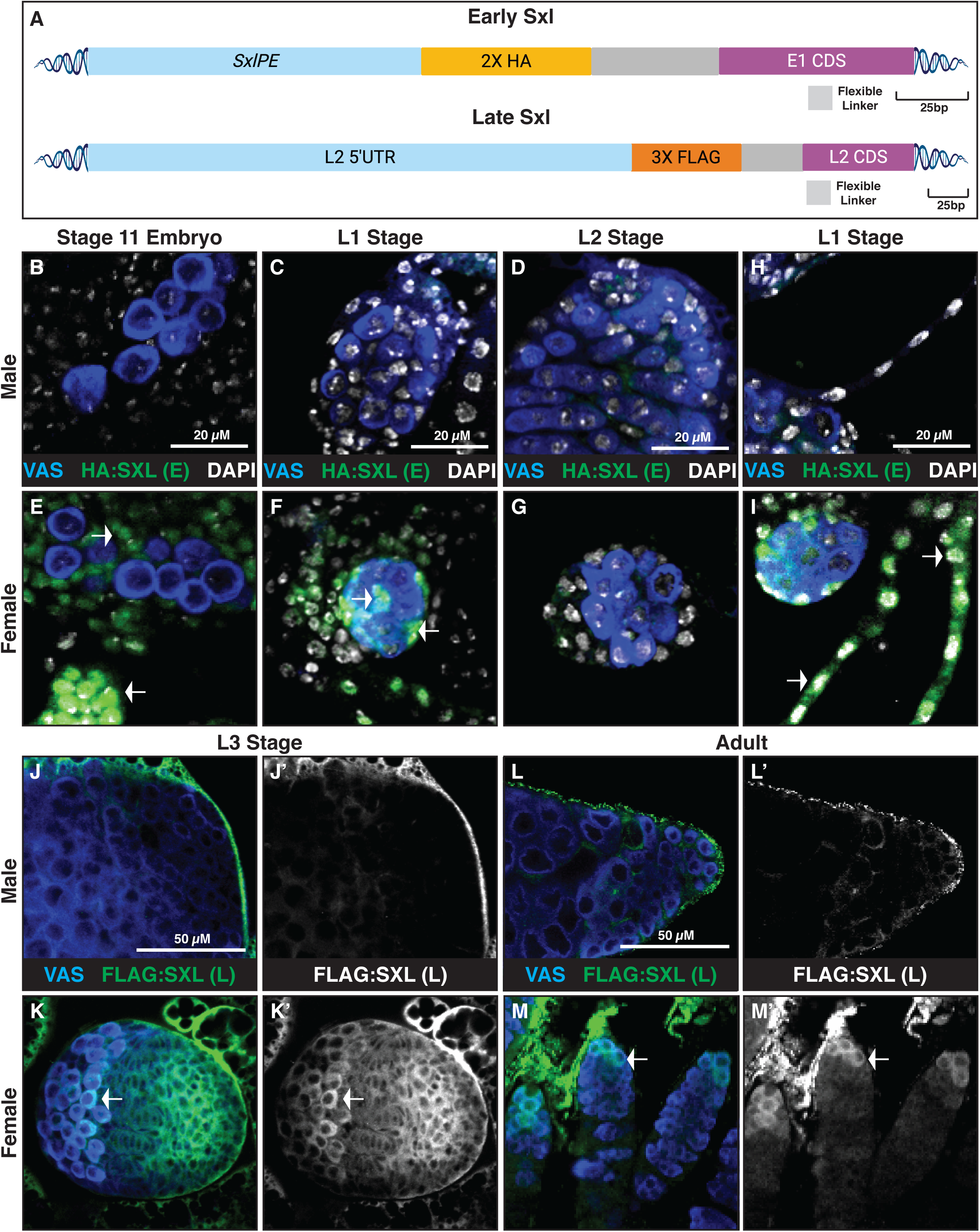
Endogenous tagging of Early and Late Sxl isoforms. A) Cartoon showing relevant portion of the *Sxl* gene locus and location of endogenous tags for Early and Late Sxl isoforms. A 2X HA tag is inserted at the N-terminus of Exon E1’s CDS to generate the HA:SxlE1 ‘Early (E) Sxl’ tag. A 3X FLAG tag is inserted at the N-terminus of Exon L2’s CDS to generate the FLAG:SxlL2 ‘Late (L) Sxl’ tag. Locus is drawn to scale. Tags are not drawn to scale. CDS: Coding sequences. UTR: Untranslated Region. B-G) Immunofluorescence of developing gonads to characterize HA:SxlE1 expression from stage 11 of embryogenesis to the second larval instar stage (L2). Note the presence of sex-specific Early Sxl expression in the nuclei of female somatic cells. Arrows mark HA-positive somatic cells. H-I) Immunofluorescence of developing guts to character HA:SxlE1 expression at the first larval instar stage (L1). Note the presence of sex-specific Early Sxl expression in the nuclei of female gut cells. Arrows mark HA-positive gut cells. J-M’) Immunofluorescence of developing gonads to characterize FLAG:SxlL2 expression from the third larval instar stage (L3) to adult stage. Note the presence of sex-specific Late Sxl expression in the cytoplasm of female germ cells. Arrows mark FLAG-positive germ cells. VAS stains germ cells. DAPI stains DNA (nucleus).

**Figure S4:**
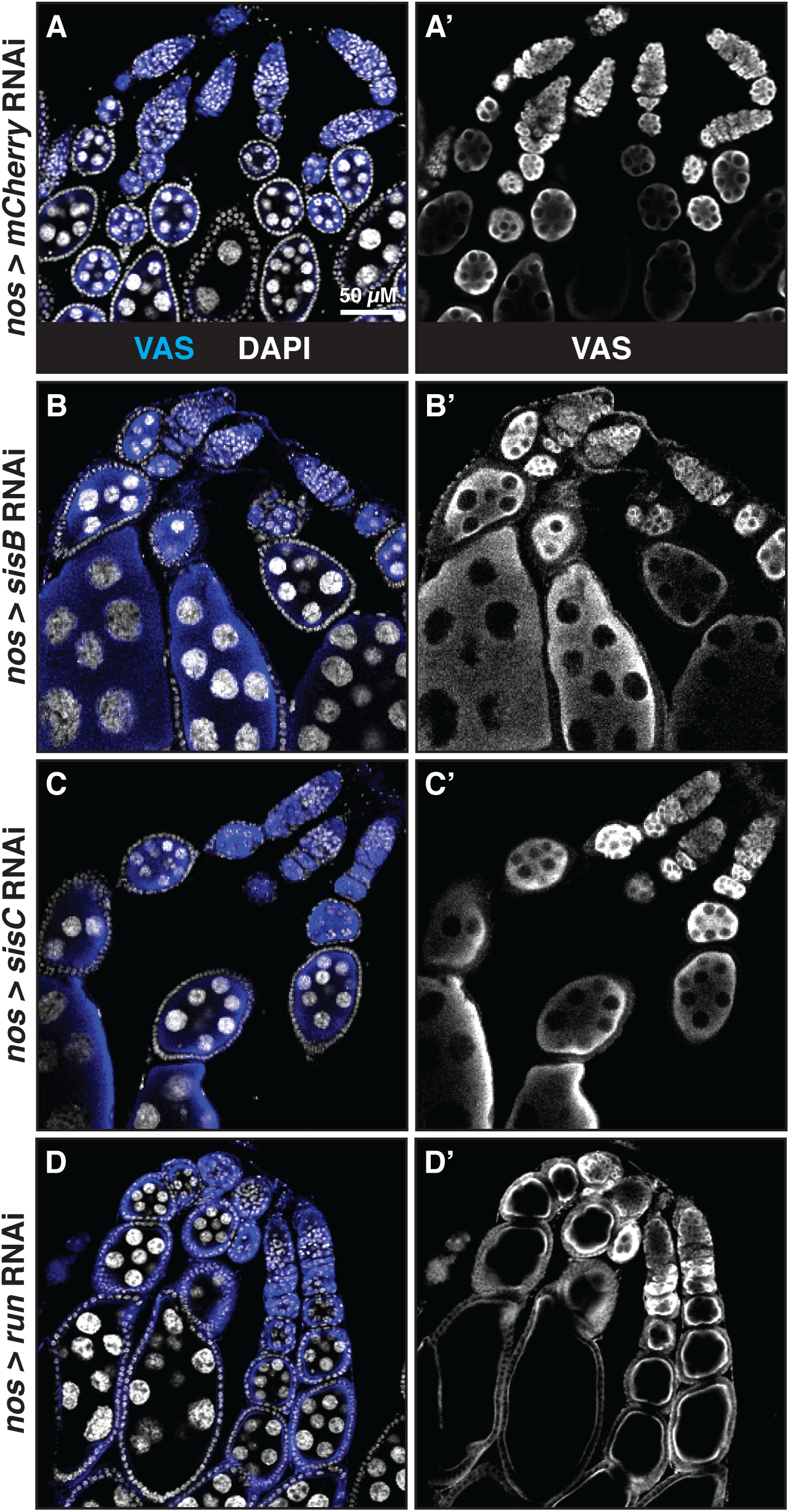
RNAi against most somatic XSEs does not affect the female germline. A-D ’) Immunofluorescence of adult ovaries to characterize germ cell phenotypes resulting from RNAi against somatic XSEs. A-A’) Wildtype ovary from fly with germline-specific *mCherry* RNAi. B-B’) Ovary from fly with germline-specific *sisB* RNAi. C-C’) Ovary from fly with germline-specific *sisC* RNAi. D-D’) Ovary from fly with germline-specific *run* RNAi. All ovaries resemble the wildtype control. VAS stains germ cells. DAPI stains DNA (nucleus).

**Figure S5:**
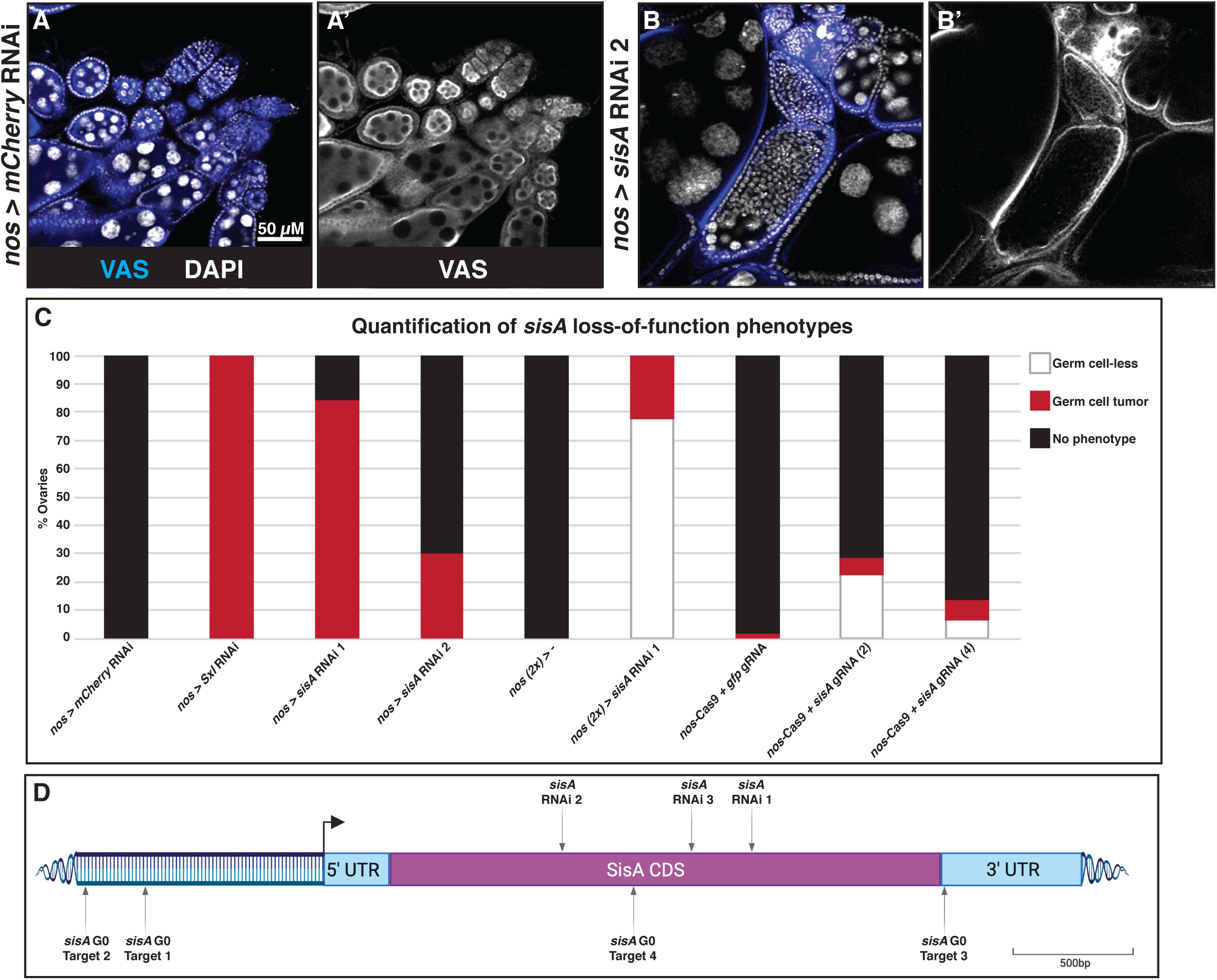
*sisA* loss of function in the female germline. A-B’) Immunofluorescence of adult ovaries to characterize germ cell phenotypes. A-A’) Wildtype ovary from fly with germline-specific *mCherry* RNAi. B-B’) Ovary with germ cell tumors from fly with germline-specific *sisA* RNAi (*sisA* RNAi 2). Note that the severity of tumors is less than those observed with *sisA* RNAi 1. VAS stains germ cells. DAPI stains DNA (nucleus). C) Graph showing percentage of ovaries exhibiting either wildtype, germ cell tumor, or germ cell-less phenotypes resultant from different loss of function conditions. D) Cartoon showing extended *sisA* locus with target sites for RNAi and guide RNA targets for G0 CRISPR. Locus is drawn to scale. CDS: Coding sequences. UTR: Untranslated Region.

**Figure S6:**
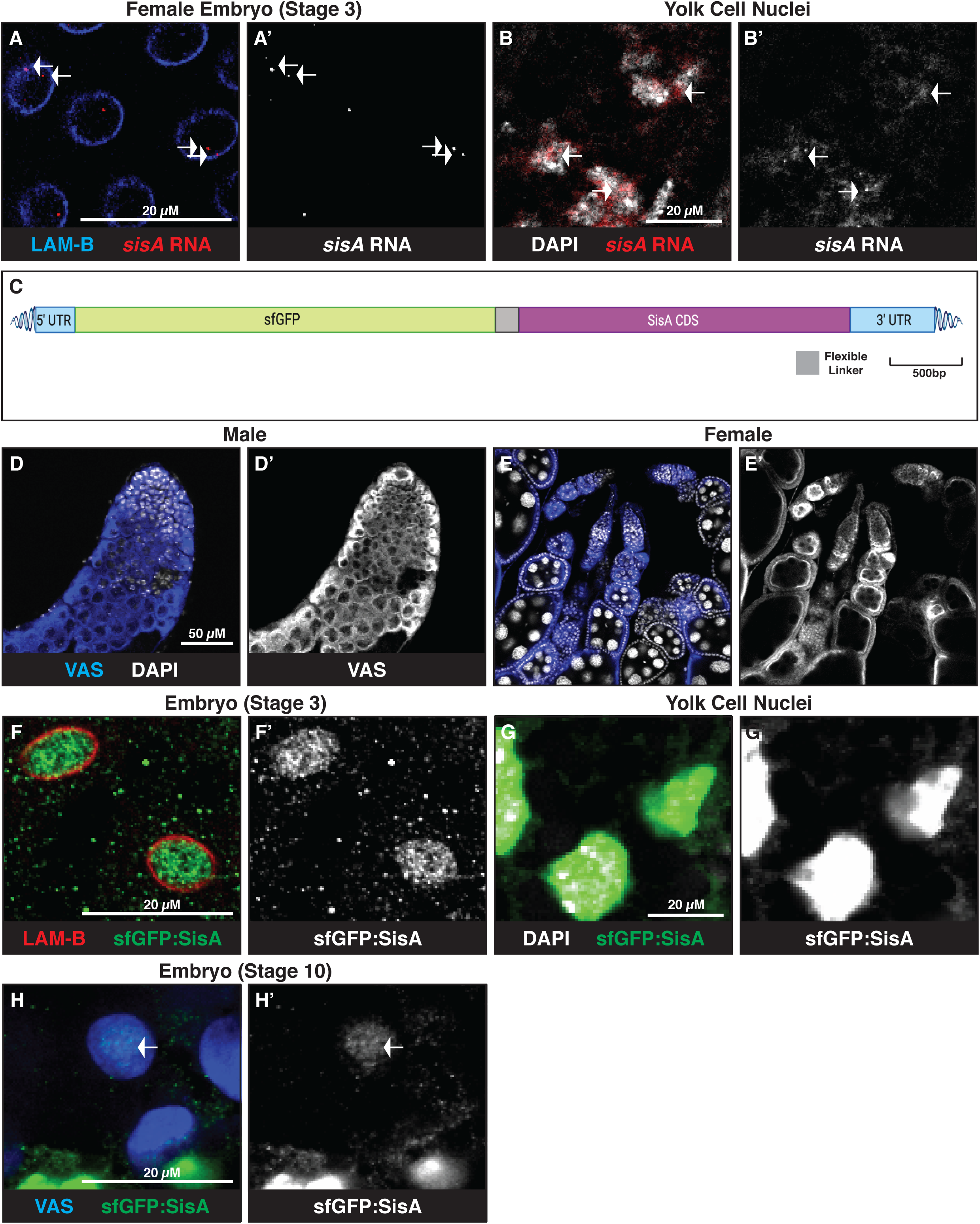
Characterizing *sisA* expression. A-B’) RNA-FISH (+ Immunofluorescence) against *sisA* in embryonic somatic cells. A-A’) Female somatic cells at stage 3 with *sisA* RNA-FISH signals. Note that two foci are observed per nucleus indicating the presence of two X chromosomes that are unpaired. Arrows mark fluorescent nuclear foci of RNA-FISH signals. B-B’) Yolk cell nuclei with *sisA* RNA-FISH signals. Arrows mark fluorescent nuclear foci of RNA-FISH signals. DAPI stains DNA (nucleus). C) Cartoon showing the *sisA* gene locus and location of endogenous sfGFP tag. An sfGFP tag is inserted at the N-terminus of *sisA*’s CDS to generate the sfGFP:SisA tag. Locus is drawn to scale. Tag is not drawn to scale. CDS: Coding sequences. UTR: Untranslated Region. D-E’) Immunofluorescence of adult gonads to characterize germ cell phenotypes caused by the N-terminal sfGFP tag on SisA. The gonads resemble wildtype gonads, suggesting that the tag does not impair SisA function. F-F’) Immunofluorescence of female somatic cells at stage 3 expressing sfGFP:SisA. G-G’) Immunofluorescence of yolk cell nuclei expressing sfGFP:SisA. Note that this expression is nuclear, consistent with SisA’s characterization as a bZIP transcription factor. H-H’) Immunofluorescence of embryonic stage 10 PGCs from flies bearing sfGFP:SisA (SisA tag). LAM-B stains nuclear lamina. VAS stains germ cells. DAPI stains DNA (nucleus).

**Figure S7:**
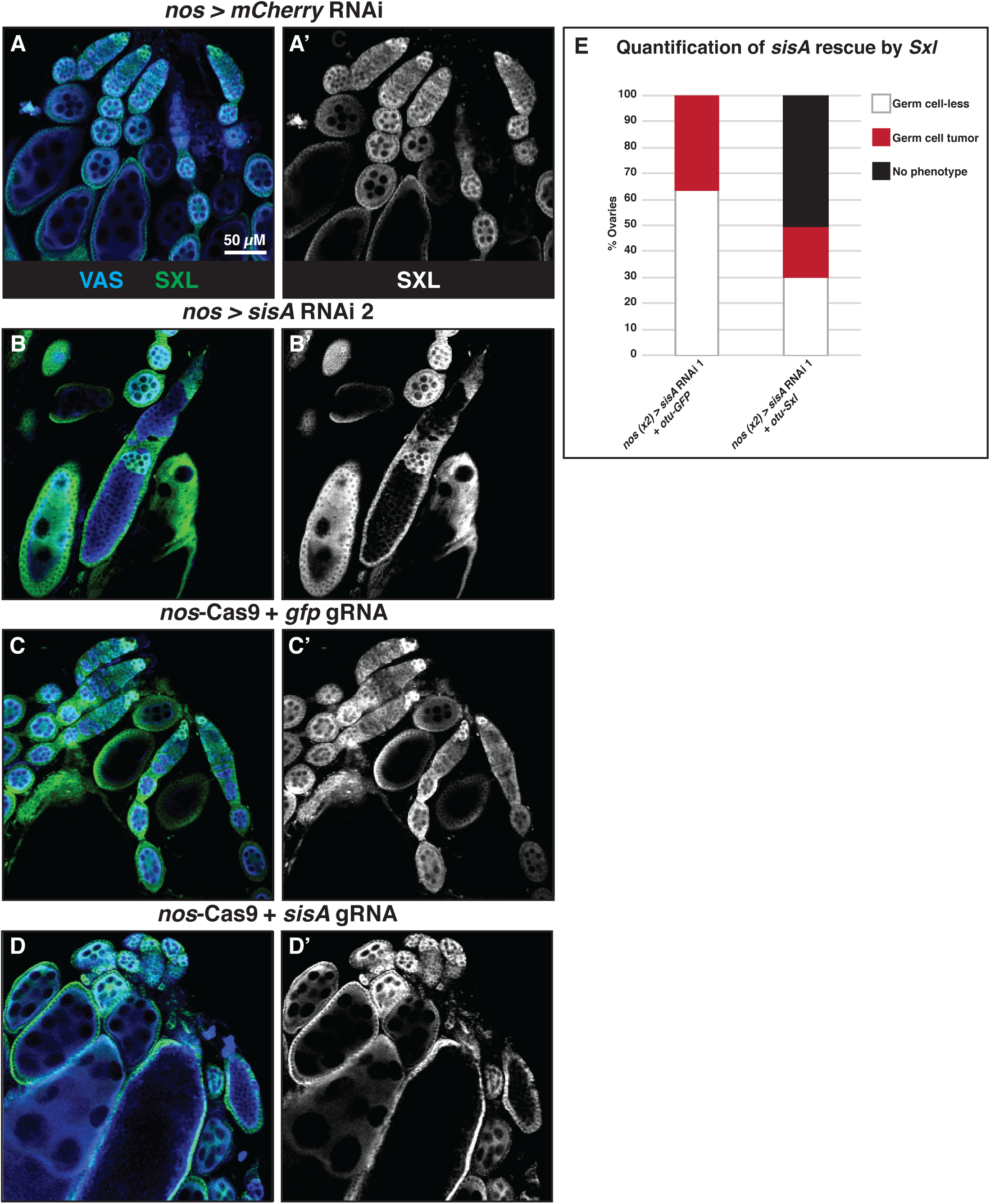
*sisA* lies upstream of *Sxl* in the female germline. A-D ’) Immunofluorescence of adult ovaries to characterize Sxl expression. A-A’) Wildtype ovary from fly with germline-specific *mCherry* RNAi. Sxl expression is highest in the early germ cells and decreases in differentiating germ cells. B-B’) Ovary with germ cell tumors from fly with germline-specific *sisA* RNAi (*sisA* RNAi 2). Note that tumorous germ cells lack Sxl expression. Somatic Sxl remains unaffected. C-C’) Wildtype ovary from fly with mutations in the germline against *gfp*. Sxl expression is wildtype. D-D’) Ovary with germ cell tumors from fly with mutations in the germline against *sisA*. Note that tumorous germ cells lack Sxl expression. VAS stains germ cells. SXL stains Sxl. E) Graph showing percentage of ovaries exhibiting either wildtype, germ cell tumor, or germ cell-less phenotypes, with or without rescue of *sisA* loss of function by *Sxl*.

Cartoons in Figures S1A, S2A, S3A, S5D, and S6C were created with Biorender.com

